# A database of plant heat tolerances and methodological matters

**DOI:** 10.1101/2025.09.14.676187

**Authors:** Timothy M Perez, Alyssa Kullberg, Evan Rehm, Kenneth Feeley

## Abstract

**Motivation:** Plant heat tolerance data are increasingly valued for their potential to help increase our understanding of species’ responses to extreme temperatures, but these efforts are hindered by methodological inconsistencies and missing contextual information. To address this issue, we collated data that compiles heat tolerance estimates and documents key sources of variation attributable to taxonomy, methodology, geography, and cultivation to improve data clarity and usability. This resource is designed to catalyze more rigorous and ecologically meaningful syntheses by enabling researchers to identify, account for, and test the drivers of variation in plant heat tolerances and their consequences.

**Main types of variable contained:** Heat tolerance estimated in degrees Celsius from photosynthetic tissue

**Spatial location and grain:** Global in scope with undersaturated taxonomic sampling and underrepresented geographic regions.

**Time period and grain:** 1935-2024

**Major taxa and level of measurement:** Primarily vascular plants encompassing >1700 taxa, >1000 genera and >200 families.

**Software format:** Comma-separated values

## Introduction

Given the fundamental importance of photosynthesis in plant function, photosynthetic tissues are commonly used to assess plant heat tolerance. Plant heat tolerances are often reported as the temperatures that cause significant reductions in normal biological or physiological functions, and have potential applications in breeding heat tolerant crops (Bita & Gerats, 2013; Langridge & Reynolds, 2021), and natural resources management (Allen *et al*., 2010; Rehfeldt *et al*., 2014). In ecological contexts plant heat tolerances are often touted for their potential to understand and predict plant responses to extreme temperatures, but their relevance to higher-order processes like growth, demographic rates, species distributions, or carbon sequestration remains uncertain.

The inability to draw mechanistic ecological conclusions or predict organismal vulnerability to thermal extremes from heat tolerances is partly due to a lack of standardized methods (Geange *et al*., 2021). Although assays of plant heat tolerances generally include similar procedures, differences within these steps can cause variation in heat tolerance estimates (Perez *et al*., 2021; Hauck *et al*., 2025). Variation in these steps has long been known to make direct comparisons of heat tolerances difficult among studies (Lange, 1965).

Different physiological responses used to measure heat tolerances with otherwise identical procedures also result in contrasting heat tolerance estimates and the inferences they generate. For example, cell vitality, cell membrane stability, and photosystem II (PSII) function are physiological traits commonly assayed in leaves and other photosynthetic tissues to estimate heat tolerance. Cell death indicates leaf tissue necrosis that is generally quantified by visual inspection including staining, microscopy techniques, and estimating the ratio of dead to living tissue (e.g. Onwueme, 1979; Chen *et al*., 1982; Nobel, 1984; Larcher *et al*., 1989). High temperatures that cause cell and organelle membrane damage can be measured with electrolyte leakage. The leakage of cellular contents is typically measured using changes in electrical conductivity. Chlorophyll *a* fluorescence may also signal the disruption of thylakoid membranes in chloroplast caused by high temperatures (Schreiber *et al*., 1976; Wahid *et al*., 2007; Zhu *et al*., 2024). However, chlorophyll *a* fluorescence is commonly used to assess processes closely related to photosystem II (PSII) function, which is temperature sensitive (Krause & Santarius, 1975; Baker, 2008).

Fluorescence from the PSII complex, measured using a minimal quantity of light (F0, i.e. fo), can increase as photosynthetic tissues are heated, signaling the closure of PSII reaction centers. A minimum pulse of light is used to obtain a value of F0, while a saturating pulse of light is used to completely reduce all PSII’s reaction centers and cause a peak in fluorescence termed the maximum fluorescence (Fm). The difference between Fm and F0 is termed variable fluorescence (Fv). Fv/Fm (FvFm) provides an estimate of the maximum quantum efficiency of PSII. Impairment of PSII function is commonly measured using relative changes in F0 or FvFm parameters.

These traits reveal physiological information at increasingly finer levels of cellular function while decreasing in biological complexity. For example, the loss of PSII function doesn’t necessarily indicate loss of cell vitality or membrane integrity. Conversely, the function of cell membranes and PSII have little physiological relevance at cell death. Consequently, these heat tolerances represent different physiological processes and should not be considered interchangeable.

The methodological inconsistencies and physiological nuances among studies are inconspicuous, but are important considerations for avoiding misuse of or misinterpretations from heat tolerance data. For example, despite limited evidence, the heat tolerance of photosystem II has been conflated with the temperature causing leaf death and has been used to make predictions about global forest mortality and ecosystem tipping points (Doughty *et al*., 2023; Winter *et al*., 2024) Similarly, some studies have compiled heat tolerance estimates from differing physiological processes - e.g., electrolyte leakage and photosystem II function (Araújo *et al*., 2013) or respiration and carbon assimilation (Lancaster & Humphreys, 2020) - to infer broad patterns in plant evolution and vulnerability to climate change. However, these different physiological mechanisms confound clear inferences regarding broader ecological and evolutionary processes.

Clarifying the methods used to estimate heat tolerance is essential for drawing meaningful conclusions about underlying physiological mechanisms and for linking them to broader phenomena such as organismal performance. We compiled a database of plant heat tolerance estimates for photosynthetic tissues and include other variables to disambiguate several major sources of variation among studies. Our database includes 1) taxonomic, 2) methodological, 3) biogeographical, and 4) cultivation data as a resource for researchers. We describe the data we collated in our database, summarize it, and briefly highlight the effects that methodological variation can have on heat tolerance estimates. Our goal is to catalyze heat tolerance research that links physiological to higher-order ecological and evolutionary processes.

## Methods

### 1) Data acquisition

To assemble our database, we identified peer-reviewed articles following forward and backward citation procedures and cross-referencing other datasets that report heat tolerance for studies published up until year 2024. Forward and backward citation procedures are an established method particularly useful for collecting empirical data (Foo et al 2021). We collected data from articles located through our citation searches if they contained heat tolerances that were 1) reported for terrestrial plants, 2) in units of temperature, 3) for photosynthetic tissues, and 4) did not quantify carbon assimilation or respiration which have databases treated elsewhere (e.g. Kumarathunge *et al*., 2019; Atkins et al). A flowchart illustrating the main steps we used to determine the eligibility of data to include is provided in our Supporting Information Section 1.

The majority of articles we collected data from were located by Perez 2019 dataset. This dataset collected by Perez 2019 was established through forward and backward citation searches over the course of dissertation research, and contributed 69 references corresponding to 941 species and 1396 heat tolerance records (Table 1). This dataset was then cross-referenced against the GlobTherm (Bennett et al 2018) and Lancaster and Humphreys (2020) for articles not currently in our dataset.

**Table 1:** Dataset summaries.

| Dataset | Dataset summary |  |  |
| --- | --- | --- | --- |
|  | References (n) | Unique species (n) | Records (n) |
| Perez, 2019 (baseline) | 69 | 941 | 1,435 |
| Perez et al. 2025 (baseline) | 131 | 1,768 | 3,197 |
| Lancaster & Humphreys, 2020 (Tmax) | 37 | 689 | 964 |
| GlobTherm (Streptophyta + Tmax ) | 12 | 73 | 73 |

*Note:*
Lancaster & Humphreys, 2020 restricted to Tmax rows. GlobTherm restricted to Streptophyta with Tmax.

Articles not already integrated into the Perez 2019 dataset were screened for compatibility. To ensure the consistency and fidelity of data in our database, we collected metadata directly from compatible articles instead of these pre-existing datasets. This was necessary since data from existing datasets may have obfuscated physiological metrics that we reported, excluded some metrics of heat tolerance, and often omitted data for species in the article they referenced. A diagram illustrating how references from other databases were incorporated into our database and overlap with one another is provided in Section 1 of our supplemental materials. A list of the data sources is found in the Supplemental Information *Table S1.* Our description of the heat tolerance metadata we collected and our procedure for collecting it described in the next section.

### 2) Taxonomic Data & Description

Taxonomic data was recorded for each heat tolerance estimate in our database (see 2.1). We updated taxonomic data (see 2.2) from the original sources to account for any changes in nomenclature.

### 2.1 original_species

The original genus and species name from the source of the heat tolerance estimate. Some of the reported taxonomic classifications in the literature are outdated or contain orthographic errors.

### 2.2 updated_species

To standardize the taxonomic information in the database, we used the ‘wcvp_match_names’ function in the ‘rWCVP’ R package (Brown *et al*., 2023) to update the *original_species* data according to the World Checklist of Vascular Plants (Govaerts et al 2021). Then we used the ‘resolve_multi’ function to select the best fitting name when multiple matches were present for a single supplied taxon. Lastly, we inspected and corrected all rows where the final match was not exact, and manual corrections were made. Genus and family names were entered when higher-resolution taxonomic data were not evident.

### 2.3 wcvp_authors

The taxonomic authorities for a given species recorded in the *updated_species* data and according to the WCVP database.

### 2.4 family

Indicates the plant family associated with the *updated_species* names.

### 3) Methodological Data & Description

We collected methodological data associated with each estimate of heat tolerance (*3.1*). This methodological data includes the type of heating method (3.2), duration of heat treatment (3.3.1), length of recovery time following heat treatment (3.4.1), the physiological method used to measure the response to heat treatment (235), and the metric used to estimate heat tolerance (3.6.1). To simplify the methodological variation across studies, we categorized the original duration of heat treatment (3.3.2), recovery time following heat treatment (3.4.2), and the original term used for each heat tolerance (3.6.2).

### 3.1 HT

We recorded the maximum heat tolerance metric that was reported per species per location within each article and reported it in the *HT* column. Heat tolerance estimates were obtained from text, supplemental data, or digitized from figures and are recorded in degrees Celsius (°C). Digitized data was obtained using the plot digitizer web app (https://plotdigitizer.com/about).

### 3.2 static_dynamic

We recorded the type of heating method used to study plant heat tolerances.

Values are either “static” or “dynamic”. Static refers to heat treatments that expose tissues to a given temperature for a fixed period of time. Dynamic refers to heat treatments that expose tissues to temperatures that change at a constant rate, often raising temperature to a high-temperature target over a given period of time (Lutterschmidt & Hutchison, 1997). Both static and dynamic heat methods may use a variety of heat sources (e.g., peltier devices, water baths, heating plates), but these are not provided in our database.

#### 3.3.1 original_duration

We recorded the duration of heat treatment used to estimate heat tolerances in minutes. Evidence suggests that the duration of heat treatment influences estimates of heat tolerances (e.g. Columbo and Timmer 1992, Cook *et al*., 2024). More specifically, increasing the duration of heat treatment is expected to cause decreases in the estimates of heat tolerances.

#### 3.3.2 duration_category

We categorized heat treatment duration data as follows: > 0 and ≤ 5 minutes is abbreviated as (0-5]; > 5 and ≤ 25 minutes is abbreviated as (5-25]; > 25 and ≤ 60 minutes is abbreviated as (25-60]; > 60 minutes or more is abbreviated as (60-inf]; and “NA” is recorded if no duration was specified. These intervals were selected to distribute heat tolerances into groups that would facilitate comparisons that are also easily implemented by practitioners.

#### 3.4.1 original_recovery

We recorded recovery time as the minutes elapsed following removal of tissues from heat treatment until the damage was quantified. Recovery time can influence heat tolerances because some physiological functions can recover following heat treatments (Havaux, 1993; Kitao *et al*., 2000; Krause *et al*., 2010). Recovery times were recorded as zero for heat tolerance estimates determined with dynamic heating methods since in these cases the damage is assessed simultaneously to treatment.

#### 3.4.2 recovery_category

We categorized heat treatment recovery times from the original_recovery data as follows: 0 and ≤ 1 minute is abbreviated as [0-1]; > 1 and ≤ 15 minutes is abbreviated as (1-15]; > 15 and ≤ 720 minutes is abbreviated as (15-720]; > 720 and ≤ 1440 minutes is abbreviated as (720-1440]; > 1440 and ≤ 2880 minutes is abbreviated as (1440-2880]; > 2880 minutes or more is abbreviated as (2880-inf]; and “NA” is recorded if no recovery time was specified. These intervals were selected to distribute heat tolerances into groups that would facilitate comparisons that are also easily implemented by practitioners.

### 3.5 method

Different physiological response variables may be measured to assess damage following heat treatments. We recorded the physiological response variable associated for each heat tolerance estimate in the method column of our database. We collected heat tolerance data and recorded any associated methods as long as they met the criteria described above.

#### 3.6.1 original_term

Once tissue damage is quantified it can be used to estimate the temperature that causes a predefined level of physiological impairment. There are several different metrics of heat tolerances that have been reported, but these tend to correspond to the temperatures that cause the first signs of impairment, cause a 50% change in the physiological response variable, or cause a near complete change in the physiological response variable. We recorded the original term for the heat tolerance from the source article in the metric columns of our database.

#### 3.6.2 HT_standardized

The original terms reported from studies were categorized as Tcrit if the reported metric calculated represents the lowest temperature reported that leads to an initial measurable or significant reduction in the physiological response measured following heat treatment, T50 if the metric calculated represented a 50% change in the physiological response measured following heat treatment, or Tmax if the metric calculated represented a maximal reduction in function of the physiological response variable being measured. The standardized metrics are recorded in the HT Standardized column of our database. Importantly, these categories are meant to refer to heat-induced *change* in a given biophysical response variable relative to control or ambient conditions, and are independent from other methodological considerations.

### 4) Biogeographical Data

Geography may influence heat tolerances through effects of climate, microclimate (Feeley *et al*., 2020; Perez, 2020; Perez & Feeley, 2020), acclimation, or local adaptation (Knight & Ackerly, 2003; Zhu *et al*., 2018). We recorded provenance and growth location information when possible in an attempt to partially account for these effects. Provenance data indicated the location that the plant tissues originated (see 4.1 & 4.2), and growth location indicated where plants were grown before tissue were sampled for measurement (see 4.3 & 4.4). We also recorded if plant tissues were grown in outdoor environments or controlled environments in our cultivation data (section 4).

We recorded the coordinates for provenance or growth sites when they were reported. These coordinates were reviewed and altered for more accuracy when necessary. For example, coordinates that were rounded at the time of reporting leading to inaccuracies (i.e. occurring in water) were altered when additional information was present that would allow more precise geolocation. We used Google Maps (https://www.google.com/maps) and GEOLocate webapps (https://www.geo-locate.org/web/WebGeoref.aspx) to determine approximate coordinates of definitive locations or landmarks when they were provided but geographic coordinates were not. When only vague general locations were provided, we inferred probable approximate coordinates using the information provided from source articles and accessible information like vegetation cover, elevation, and our expert judgement. If we could not confidently determine the geolocation within 10km of its most likely ecosystem collection location, no coordinate was recorded.

We mapped the growth site locations for all records that also reported sample provenance data to the Intergovernmental Panel on Climate Change (IPCC) Working Group I Reference Regions and Whittaker biome space to characterize the biogeographical distribution of the data represented within our database. The IPCC regions represent sub-continental areas used for broad-scale biophysical climate modeling in the Coupled Model Intercomparison Project (Iturbide *et al*., 2020), and Whittaker biomes characterize biomes based on coarse temperature and precipitation gradients. Data were mapped to the Whittaker biomes based on the *plotbiomes* R package (Stefan and Levin (2018).

### 4.1 provenance_latitude

These data record the latitude in decimal degrees where plant material is originally from prior to heat tolerance measurement.

### 4.2 provenance_longitude

These data record the longitude in decimal degrees where plant material is originally from prior to heat tolerance measurement.

### 4.3 growth_site_lat

These data record the latitude in decimal degrees where plants were grown prior to heat tolerance measurement.

### 4.4 growth_site_lon

These data record the longitude in decimal degrees where plants were grown prior to heat tolerance measurement.

### 4.5 location_name

These data record the location that the heat tolerance measurements were made and used to gather provenance or grow site information where applicable.

### 5) Cultivation Data

Our database focuses on undomesticated plant species but does contain data for some domesticated and cultivated species. The distinction between wild and domesticated or cultivated species is likely relevant to researchers with differing expertise, and may be useful for understanding the effects of selective breeding on heat tolerance. We recorded cultivation status of each species (5.1) and basic information on the growing conditions of the plants used to estimate heat tolerances where possible (5.2).

### 5.1 Agricultural_sp

Records the cultivated and or agricultural status of species measured.

Species were considered agricultural or cultivated if they are widely seen as commercial crops, commodities, crops grown for sustenance, or ornamental purposes. In some cases, the original reference provided information to help determine if the specific individuals used were of cultivated varieties. We cross-referenced all species in our database against the Crop Wild Relative (CRW) database, which contains scientific names of cultivated plants (Crop Wild Relative Occurrence data consortia, 2024). Values are recorded as “Y” or “N” indicating that yes, or no, respectively.

### 5.2 growing_conditions

This data categorizes the growing conditions the plants were grown in prior to heat tolerance assessment. These data include greenhouse, common garden, growth chamber, or *in situ* designations. Importantly, we do not include information about soil, water or temperature, or light regimes, which can all influence heat tolerances.

### 6) Bibliometric Data

We provide full citations for each study from which heat tolerance estimates and their associated metadata were obtained (Section 6.1).

### 6.1 reference

We recorded the authors, publication date, study title, and journal or analogous information for each heat tolerance estimate.

### 7) Correlations among methods

We conducted pairwise Pearson correlation analyses between all combinations of heat tolerance measurement methods to understand which physiological responses exhibit coordinated temperature thresholds. For each pair of methods, we identified overlapping species and averaged replicate measurements within species for each method. Correlations were calculated only when at least five species were shared between methods. We recorded the Pearson correlation coefficient (r), associated p-value, and the number of overlapping species (n) for each method pair. These values were compiled into matrices representing correlation strength, significance, and sample size, respectively.

## Results

### 1) Data acquisition

Our final dataset contained 3197 heat tolerance records which is three times as many records as the Lancaster and Humphreys dataset, and over 43 times the number of records for terrestrial plants as the Globtherm dataset (Table 1). Our heat tolerance records correspond to 1745 taxonomic entities, which is over twice as many species as the Lancaster and Humphrey (2020) dataset, and over 23 times the number of records for terrestrial plants in the Globtherm dataset (Table 1). Notably, our database contains 91 unique references corresponding to 1231 species records and 2177 records of heat tolerances not present in other datasets.

Despite sharing references, our data collection procedure led to considerable differences in the number of species and records in our final database compared to those of GlobTherm and Lancaster and Humphreys. For example, of the 11 references common to both datasets, 72 species and 72 records were reported from GlobTherm while we reported 93 species and 239 records (Table 2). Lancaster and Humphreys reported 637 species and 856 records while we reported 622 species and 905 records for the same 30 references (Table 2). Reference-level explanations for differences in the number of species and heat tolerance records between our database and the GlobTherm and Lancaster and Humphreys datasets are provided in Supplemental Tables 1 and 2.

**Table 2:** Pairwise overlaps between datasets (counts within each overlap).

| Pair | Dataset | Overlap summary |  |  |
| --- | --- | --- | --- | --- |
|  |  | References (n) | Unique species (n) | Records (n) |
| Lancaster & Humphreys, 2020 ↔ Perez, 2019 | Lancaster & Humphreys, 2020 (overlap with Perez, 2019 | 16 | 566 | 716 |
|  | Perez, 2019 (overlap with LH) | 16 | 545 | 778 |
| GlobTherm (Streptophyta + Tmax) ↔ Perez, 2019 | GlobTherm (Streptophyta + Tmax overlap with Perez, 2019 | 7 | 63 | 63 |
|  | Perez, 2019 (overlap with GlobTherm) | 7 | 81 | 222 |
| Lancaster & Humphreys, 2020 ↔ Perez et al. 2025 | Lancaster & Humphreys, 2020 (overlap with Perez et al. 2025 | 30 | 637 | 856 |
|  | Perez et al. 2025 (overlap with LH) | 30 | 622 | 906 |
| GlobTherm (Streptophyta + Tmax) ↔ Perez et al. 2025 | GlobTherm (Streptophyta + Tmax overlap with Perez et al. 2025 | 11 | 72 | 72 |
|  | Perez et al. 2025 (overlap with GlobTherm) | 11 | 93 | 239 |

Note:
Counts reflect the subset defined by shared references in each pair. Lancaster & Humphreys, 2020 restricted to Tmax rows. GlobTherm restricted to Streptophyta with Tmax.

### 2) Taxonomic Data

Our database represents heat tolerance estimates belonging to 1768 different taxa classified at the species, genus or family levels. These data correspond to 1001 genera and 250 plant families. The heat tolerances for the 50 most commonly reported families and genera are depicted in Figure 1 (D & E). The species in our database represent predominantly vascular plants, but include some moss species. A stable version of the database present in this manuscript is stored in (http://datadryad.org/share/LINK_NOT_FOR_PUBLICATION/k7Zb6yQwimY9E84FZQ4NILE1j7SJK8vraLRsVG1zyn8). Updated versions of this database will be hosted on (authors github project website).

**Figure 1:**
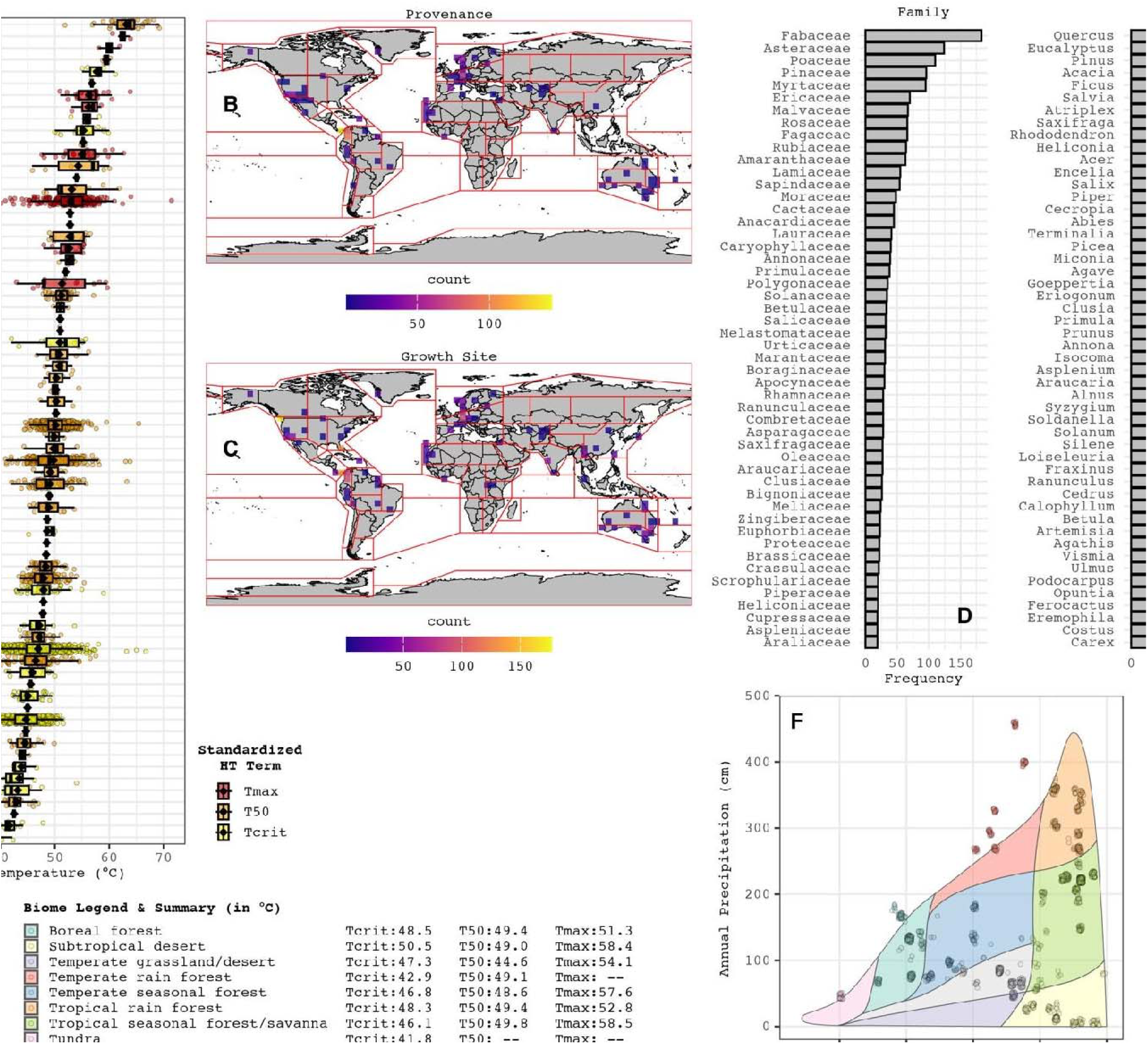
(A) Boxplot of 3197 heat tolerance estimate colored by standardized HT metric (i.e. Tcrit, T50, or Tmax) across 70 distinct methods ordered by mean (black diamonds). Numbers on y-axis correspond to distinct method reported in Supplemental Table X); Species sample size within each 500×500km pixel with provenance (B) or growth site (C) coordinates for each IPCC region (pixel color indicates). The 50 most commonly reported plant families (D) and genera in the database; (F) Biome representation of records in our database. Points represent heat tolerances records mapped to growth site climates after filtering for individuals grown *in situ* or common gardens with known provenance data (points jittered to illustrate sample sizes).

### 3) Methodological Data

We recorded dynamic versus static heating methods, heat treatment duration, recovery durations, physiological response variables, and the term used to report each heat tolerance. The original reported metadata associated with each heat tolerance represented 139 distinct methods and terms among 131 studies we used to gather data. Our categorizations and standardization procedure resulted in 70 distinct methods. Data from these standardized methods are depicted in Figure 1A with a total number of records associated with each in Supplemental Table 4). We noted that 59 different terms used to describe the heat tolerances we categorized into one of three metrics (Table S3), that 17 methods represented by a single record of heat tolerance.

### 3.1 HT

Our database contains heat tolerances that are summarized in Figure 1A. Descriptive statistics associated with Figure 1A are included in Supplemental Table 4, which includes the full complement of methodological metadata for each numbered category. For example, group 1 with the highest mean heat tolerance of 63.1°C, and represents species that were statically heated for durations of 25-60 minutes, allowed to recover for 720-1440 minutes, before having cell death quantified and used to calculate the T50 metric.

### 3.2 static_dynamic

Our database consists of 1980 heat tolerance records estimated using static heating methods and 1213 records estimated using dynamic heating methods.

### 3.3 original_duration & duration_category

We recorded 19 different heating durations reported across all studies before categorizing them into 4 groups. We placed 1246 records into the (0-5] minute duration category, 740 records in the (5-25] minute category, 1152 records in the (25-60] minute category and 27 records in the (60-inf] minute category. Of the studies we reviewed, heat treatment durations ranged from a few seconds in dynamic heating rates (O’Sullivan *et al*. 2017) and up to 12 hours (Mooney & Billings 1961), but durations >60 minutes were relatively rare.

### 3.4 original_recovery & recovery_category

We recorded 25 different original recovery times reported among studies and categorized them into 6 groups. There were 1273 different records for the [0-1] minute recovery time category, 11 records for the (1-15] minute recovery time category, 126 records for the (15-720] minute recovery time category, 1273 records for the (720-1440] minute recovery time category, 81 records for the (1440-2880] minute recovery time category, and 312 records for the (2880-inf] minute recovery time category.

### 3.5. *method*: We reported five different physiological methods used to estimate heat tolerance

These methods include cell death, electrolyte leakage, FvFm, fo-rise, and protoplasmic streaming. There are 475 heat tolerances assessed with cell death, 1191 assessed with fo-rise method, 1339 assessed with FvFm, 147 assessed with electrolyte leakage, and 41 that measured protoplasmic streaming. The highest heat tolerance was assessed using cell death, and the lowest were assessed using the fo-rise method.

### 3.6 original_term & HT_standardized

We compiled 59 distinct terms used to describe heat tolerance estimates and categorized them into three main types: Tmax, T50, and Tcrit (Supporting Information Table S1). We assigned 30 distinct terms to the Tcrit category, 20 different terms as T50, and 9 different terms were classified as Tmax.

### 4) Biogeographical Data

We recorded 1519 records with provenance coordinates (Figure 1B), 2913 records with growth site coordinates (Figure 1C), and 1322 heat tolerance records with provenance and growth coordinates that represented data from individuals grown *in situ* or in common garden environments. These 1322 records corresponded to 872 species from 46 sources and were used to calculate mean heat tolerances for IPCC regions and Whittaker biomes.

Estimates of Tcrit were available for 24 of the 58 IPCC regions, T50 was available for 21 regions, and Tmax was available for 11 regions (Table S2). Central-American IPCC regions were the best represented in our dataset (n=305) followed by South American (n=279), and European (N=61) regions. Over 50% of IPCC regions are not represented by any heat tolerance estimates in our database even when only provenance or growth location data are considered. Estimates of Tcrit were available for all 9 Whittaker Biomes, T50 were available for all biomes except the tundra, and at least one estimate of Tmax was available for all biomes except the tundra and temperate rainforests (Figure 1F). We reported the averages of each heat tolerance metric regardless of method in Table S5 for IPCC regions and in Figure 1F for biomes.

### 5) Cultivation Data

We collected agricultural status and growth condition metadata associated with each heat tolerance record. We classified 108 species (6% of all species in our dataset) as having agricultural or cultivated status, which accounted for 215 heat tolerance records (0.6% of all records). The four most frequently represented agricultural species were *Pisum sativum* (n = 9), *Mangifera indica* (n=6), *Prunus avium* (n=6), and *Zea mays* (n=6). The three most common agricultural plant families were Fabaceae (n = 32), Poaceae (n = 26), and Solanaceae (n = 16). Our database contains 1717 *in situ*, 973 common garden, 248 greenhouse, and 72 growth chamber growth heat tolerance records. We were unable to obtain growth conditions for 179 records.

### 6) Bibliometric Data

We collected data from 131 different studies published from 1935-2024. Four studies contributed nearly one-third of all records (33.1%) of the database records: Bison and Michaletz (2024, n = 354); Slot et al. (2021 n = 288); Perez and Feeley (2020, n = 246), and O’Sullivan et al (2017, n=168) However, 38.0% of all records in the database are from studies (n = 116) that report fewer than 50 records.

### 7) Correlations among methods

Only 58 pairwise comparisons were possible of the 2415 unique combinations (excluding self-correlations) across all methods in our database. We observed 18 statistically significant (0.05 ≥ p) correlations among 19 different methods for measuring heat tolerance (Table 3). The majority of the correlations we observed were between identical methods that differed in one procedural step or standardized term (i.e. Tcrit, T50 or Tmax), but were otherwise identical. Notable significant correlations were observed between fo-rise and cell death methods (Table 3: methods 7 & 8 and 8 & 9), electrolyte leakage and cell death methods (Table 3: 27 & 30). Many significant correlations were observed between fo-rise and FvFm methods. All 2415 pairwise correlations are provided in Supplemental Table 5.

**Table 3:** Significant correlations between heat tolerance methods with p < 0.05.

| Method 1 ID | Method 2 ID | Method 1 | Method 2 | Correlation Summary |  |  |
| --- | --- | --- | --- | --- | --- | --- |
|  |  |  |  | r | p | n |
| 1 | 22 | static (5-25] (720-1440] FvFm T50 | static (5-25] (1440-2880] FvFm T50 | **0.934** | **0.00e+00** | **43** |
| 14 | 15 | static (25-60] (1440-2880] FvFm T50 | static (25-60] (1440-2880] FvFm Tcrit | **0.908** | **0.00e+00** | **14** |
| 7 | 10 | dynamic (0-5] [0-1] fo-rise Tmax | dynamic (0-5] [0-1] fo-rise T50 | **0.905** | **0.00e+00** | **66** |
| 9 | 10 | dynamic (0-5] [0-1] fo-rise Tcrit | dynamic (0-5] [0-1] fo-rise T50 | **0.887** | **0.00e+00** | **70** |
| 54 | 55 | static (5-25] NA FvFm T50 | static (5-25] NA FvFm Tcrit | **0.856** | **1.39e-02** | **7** |
| 24 | 58 | static (5-25] (15-720] FvFm T50 | static (5-25] (15-720] FvFm Tmax | **0.810** | **0.00e+00** | **22** |
| 8 | 9 | static (25-60] NA cell death T50 | dynamic (0-5] [0-1] fo-rise Tcrit | **0.787** | **0.00e+00** | **26** |
| 7 | 32 | dynamic (0-5] [0-1] fo-rise Tmax | static (60-inf] [0-1] FvFm T50 | **0.727** | **2.10e-03** | **15** |
| 10 | 32 | dynamic (0-5] [0-1] fo-rise T50 | static (60-inf] [0-1] FvFm T50 | **0.720** | **2.50e-03** | **15** |
| 38 | 40 | static (25-60] NA FvFm T50 | static (25-60] NA FvFm Tmax | **0.707** | **3.20e-03** | **15** |
| 9 | 32 | dynamic (0-5] [0-1] fo-rise Tcrit | static (60-inf] [0-1] FvFm T50 | **0.706** | **2.20e-03** | **16** |
| 9 | 14 | dynamic (0-5] [0-1] fo-rise Tcrit | static (25-60] (1440-2880] FvFm T50 | **0.671** | **8.60e-03** | **14** |
| 27 | 30 | static (25-60] (720-1440] electrolyte leakage T50 | static (25-60] (2880-inf] cell death Tcrit | **0.623** | **2.30e-02** | **13** |
| 9 | 15 | dynamic (0-5] [0-1] fo-rise Tcrit | static (25-60] (1440-2880] FvFm Tcrit | **0.551** | **4.10e-02** | **14** |
| 7 | 8 | dynamic (0-5] [0-1] fo-rise Tmax | static (25-60] NA cell death T50 | **0.545** | **2.20e-03** | **29** |
| 1 | 33 | static (5-25] (720-1440] FvFm T50 | static (5-25] (720-1440] FvFm Tmax | **0.511** | **6.40e-03** | **27** |
| 7 | 9 | dynamic (0-5] [0-1] fo-rise Tmax | dynamic (0-5] [0-1] fo-rise Tcrit | **0.224** | **1.00e-04** | **288** |
| 1 | 29 | static (5-25] (720-1440] FvFm T50 | static (5-25] (720-1440] FvFm Tcrit | **0.154** | **5.70e-03** | **320** |

Note:
ID's are not used elsewhere in the database and are included only here for ease of comparison.

## Discussion

### 1) Data acquisition

We compiled the largest database of heat tolerances for terrestrial plants to date. Our database doubles the number of species with heat tolerance data and more than triple the total number records compared to previous datasets for terrestrial plants. Although our database builds on previous datasets, it is distinguished by detailed metadata describing measurement methods, biogeographical locations and cultivation information previously excluded.

Comparing species and heat tolerances records derived from the same references but in different datasets revealed that in many cases our procedure for collecting data resulted in more records per reference than other datasets. This is likely explained by our collection procedures and our focus on multiple different methods of measuring heat tolerances. As a consequence, our database also reported more species for shared references compared to other datasets. However, other datasets report more species and heat tolerance records when they include physiological data that we omit like carbon assimilation, respiration, or acclimation.

### 2) Taxonomic Data

Although our database is the most comprehensive compilation of plant heat tolerance estimates to date, it represents only a small fraction of global plant diversity. The 1,768 taxonomic entities included span 1,001 genera and 250 families, yet this coverage is extremely limited when compared with the estimated >350,000 vascular plant species worldwide and their families and genera (Antonelli et al 2023). The inclusion of a few moss species highlights that non-vascular plants remain especially understudied. This taxonomic bias underscores both the need for caution when extrapolating our results to all plants and the opportunity for future work to expand heat tolerance measurements across underrepresented clades and functional types.

### 3) Methodological Data

The 70 different methods in our database suggest that a novel methodological approach for measuring heat tolerances is reported from every other study we reviewed. Similarly, the 59 different terms we observed equate to more than one novel term reported for every two studies we reviewed. These results underscore the lack of standardized approaches in heat tolerance literature noted in other studies (Geange et al 2021), which may contribute to hindering the ability to link heat tolerances with higher-order ecological processes.

### 3.1 HT

### 3.2 static_dynamic

It is unclear how different methods of heating may influence physiology and resulting heat tolerances. The specific heating tools or methods used for static or dynamic heating are not recorded in this database, but commonly include hot water baths with or without direct contact with water (Colombo & Timmer, 1992), a Peltier device (e.g. Godoy *et al*., 2011), and even steam-saturated high air temperatures (e.g. Sapper, 1935). Dynamic heat treatments continuously heat leaves and are more representative of how leaves experience increases in temperature *in situ*, but confound the effect of temperature treatment and duration on heat tolerances (Krause *et al*., 2010). The effect of different heating rates on heat tolerances has not been fully investigated, but may be an important consideration for future research given leaf tissue heating rates may be influenced by a thermal time constant theorized to help modulate leaf temperature and avoid thermal damage (Leigh *et al*., 2012).

Static heating procedures do not confound the effects of temperature treatment and duration, but may not represent the cumulative effects of temperature on physiological processes. Ongoing research on the cumulative thermal stress promises a potential framework to harmonize heat tolerances estimated with static and dynamic heating approaches (Cook et al 2024).

### 3.3 original_duration & duration_category

The duration of heat treatments varies extensively in our data which may bias heat tolerance estimates. For example, the relationship between heat treatment duration and heat tolerances has been described using an exponential decay function (Gauslaa, 1984; Colombo & Timmer, 1992; Cook *et al*., 2024). This modeled relationship suggests that 30-second and 5-minute heat treatments with identical static temperature treatments can result in at least a 4.5°C difference in heat tolerance (Colombo & Timmer, 1992). Conversely, a 4.5° C difference in heat tolerance can also be observed between 20 and 160 minute durations based on this same modeled relationship. This suggests that long-duration heat tolerances may reduce the magnitude of measurement errors during heat tolerance assessments.

However, heat treatments longer than 20 minutes may allow acclimation of heat tolerances and introduce bias (Havaux, 1993). Ultimately, different species may exhibit different shapes of their exponential decay curves in response to heat treatment duration and that collating heat tolerances with different durations may introduce bias (Gauslaa, 1984; Cook *et al*., 2024). We advocate for caution when drawing conclusions from heat tolerance data aggregated from different heat treatment durations.

### 3.4 original_recovery & recovery_category

Recovery times allow for repair following heat treatment so irreversible damage can be assessed. The recovery times in our database range from non-existent - in the case of fo-rise methods representing reversible damage - to minutes and weeks in studies when measuring cell death. Different recovery times may have dramatic effects on heat tolerance estimates. For example, assessing heat damage 15 minutes and 1440 minutes (24 hours) after heat treatment resulted in a 2.3 C increase in the heat tolerances of *Ficus insipida* when assessed with FvFm chlorophyll fluorescence (Krause *et al*., 2010). Conversely, heat tolerance may decrease with recovery times when assessed using other methods like cell death (Mooney & Billings, 1961; Krause *et al*., 2015; Winter *et al*., 2024). This highlights that different physiological processes may require different recovery periods, and that recovery time reflects an underlying physiological recovery process in addition and a procedural step.

Although we report original and standardized recovery times, the recovery of the corresponding physiological processes being measured is not implicit. Ultimately, even heat tolerances of different physiological processes that are assessed with identical methods may reflect different levels of reversible or irreversible damage. Heat tolerance data with differing recovery times may indeed indicate ranges of recovery, which we do not report.

### 3.5. method

The five physiological methods of assessing heat damage we recorded were cell death, electrolyte leakage, FvFm, fo-rise, and protoplasmic streaming. These methods may largely reflect approaches used to report heat tolerances in units of temperature that we targeted. As such, our database should not be considered exhaustive of all approaches used to assess heat tolerances in photosynthetic tissues. For example, we intentionally excluded important physiological processes like carbon assimilation and respiration because they are treated elsewhere (e.g. Kumarthunge et al 2019, Atkin et al 2015). Future efforts collating disparate plant databases and physiological methods may advance heat tolerance research by revealing novel links in physiological responses.

### 3.6 original_term & HT_standardized

We categorized 54 distinct terms for heat tolerance into Tmax, T50, or Tcrit. Our standardized terms for heat tolerance are heuristics that make the excessive terminology for heat tolerances more tractable for summary and descriptive statistics. Importantly, this artificially reduces the complexity for analyzing many different methods and may compromise precision of the metrics that were originally reported.

Our database highlights the extensive and often inconsistent terminology used to describe plant heat tolerance, which partly reflects the diversity of measurement approaches. One notable exception is T50, which is consistently defined as the temperature causing a 50% change in a physiological response variable. We suspect the widespread use of T50 stems from its standardized definition based on percent change relative to baseline conditions, which makes it broadly applicable across different physiological metrics. In contrast, the terms Tcrit and Tmax are more ambiguous, often lacking a clear association with a specific threshold of physiological change or damage. Given that Tcrit and Tmax are categories of heterogeneous metrics, they may exhibit more variation when analyzed using our standardized format compared to the T50 category.

Despite extensive variation in methods and terminology, standardized approaches may not be warranted. It is possible that the lack of standardized methods reflects the lack of coordination with other physiological processes or plant performance and standardized methods may stifle innovation or novel insights. Alternatively, standardization may not be appropriate since different approaches probe distinct physiological processes. Instead, we advocate for uniform reporting terminologies of heat tolerances regardless of methods as a way to reduce the proliferation of terminology and promote comparisons among studies. For example, reporting Tcrit, T50, and Tmax heat tolerance metrics together and along with the percent damage or change associated with the Tcrit and Tmax heat tolerances is likely to facilitate interpretability of heat tolerance estimates across studies. This approach will help preserve the utility of legacy methods without discouraging novel approaches with clear physiological relevance.

### 4) Biogeographical Data

We used geolocation data to map heat tolerances to IPCC regions and Whittaker biomes (Fig 1F and Table S5). Importantly, these summaries represent a composite heat tolerance comprising many different species, methods, and physiological processes, and should be considered heuristics. The averaging we applied across estimates from varying methods obfuscates their underlying physiological data. These estimates may only be useful as a point of comparison when heat tolerance estimates for specific methods, species, and geographies are not available.

Despite the size of our database, geographic coverage of plant heat tolerances remains extremely limited. Fewer than half of the broadly defined IPCC regions are represented, and finer-scale ecoregions would show even greater gaps. This limited scope means that generalizations about global patterns in heat tolerance may be hard to achieve. This issue is exacerbated given that heat tolerances may acclimate, which means estimates from location may not be representative of those for the same species in different micro-or macro climates (Lancaster and Humphreys, 2020; Zhu et al 2018; Kullberg and Feeley, 2024). Expanding sampling efforts, particularly those that link heat tolerance to broader measures of organismal performance would likely facilitate methodological standardization and should be a global priority.

### 5) Cultivation Data

We collected data from studies that reported heat tolerances in degrees Celsius, which allows for standardized comparisons across physiological processes, species, and abiotic factors such as microhabitat or climate. This selection criteria may have biased against agricultural species that often have heat tolerances measured in units of crop yield. Since crop yield tends to be species-specific and in different units, they were incompatible with our database. Incorporating agricultural data into our database may become feasible if crop yield can be reliably linked to temperature-based measures of heat tolerance.

Our data includes growth conditions, which enable users to determine whether plants were coupled or decoupled from ambient environments. However, we did not include other variables commonly associated with growth conditions such as night-versus daytime temperatures, light regimes, or watering treatments, all of which are known or hypothesized to influence heat tolerance estimates. Incorporating these variables in future efforts may help explain additional variation in the heat tolerance data.

### 6) Bibliometric Data

Our bibliometric analysis highlights the advantages of forward and backward citation searching for assembling heat tolerance records. Our approach allowed us to systematically identify relevant empirical studies, including older or less visible work that might not appear in keyword searches. This strategy yielded a greater number of references, species, and records than comparable datasets even when we drew on overlapping sources. Nevertheless, gray literature and non-English publications remain underrepresented. However, we anticipate that broad use of this database will help surface additional sources and mitigate any biases in our collection approach.

### 7) Correlations among methods

#### Physiological limitations

The majority of methods in our database showed no correlation. However, heat tolerances estimates using fo-rise methods and electrolyte leakage were correlated with heat tolerances assessed with cell death, but not each other. Correlations among these different physiological responses suggest coordinated underlying physiological mechanisms.

These results suggest that heat tolerances for electrolyte leakage and fluorescence represent independent sub-cellular physiological processes. Conversely, many metrics of heat tolerances using fluorescence (i.e. fo-rise and FvFm) showed significant correlations, which is to be expected for closely related physiological processes. The lack of correlation between heat tolerances estimated between FvFm and cell death, may suggest physiological processes associated with FvFm may be subordinal with respect to fo-rise methods.

While cell death likely influences membrane integrity and photosynthetic function, the reverse may not be true—alterations in electrolyte leakage or chlorophyll fluorescence do not necessarily indicate imminent cell death or electrolyte leakage. Not only do these methods represent different levels of biological complexity, our results suggest they may be physiologically orthogonal given their lack of coordination. A key challenge remains understanding how different measures of heat-induced damage relate to one another and scale-up in meaningful ways to predict broader physiological or biological performance.

The lack of correlations we observed may be the result of biases inherent in the data we collected and did not account for in our database. For instance, we did not record metadata related to night or daytime growth temperatures, drought, light acclimation, or tissue and organismal ontogeny that may influence heat tolerance estimates. Studies designed to compare different methods under controlled conditions are needed to validate our results.

However, the FvFm parameter, and even less so fo, reveal few physiological insights beyond the quantum yield of PSII (Baker, 2008). The documented effects of exceeding fluorescence-based heat thresholds on broader physiological processes or organism performance is poorly documented. Cell death and electrolyte leakage supercede the sub-organelle physiological scale of fluorometrics. Heat tolerances estimated using these methods also tend to exhibit higher heat tolerances than those measured with fluorescence - possibly illustrating a logical necessity for greater tolerances at broader scales of biological organization.

## Conclusion

Plant heat tolerances are widely measured but difficult to generalize due to inconsistent methods. We compiled the largest database to date, containing over 3,000 records from over 1,700 taxa, with standardized taxonomy, methodological, geographic, and cultivation metadata. We showed that heat tolerances measured with differing physiological methods may be related, but capture different levels of biological organization and may not scale consistently to broader measures of plant performance. Despite uneven coverage across regions and lineages, our database highlights key gaps, provides a framework for disambiguating methods, evaluating their comparability, and linking heat tolerance to broader ecological and evolutionary processes.

## Author Contributions

All authors contributed equally in aspects of authorship.

## Acknowledgements

The authors declare no conflicts of interest.

## Data Accessibility Statement

All data is available from Dryad (http://datadryad.org/share/LINK_NOT_FOR_PUBLICATION/k7Zb6yQwimY9E84FZQ4NILE1j7SJK8vraLRsVG1zyn8) with updated versions located on (authors’ github here following review).

## Additional Files

***Supporting Information:***

Supporting_Information_12Sept2025

Supplemental_Table_1-2-4-6.xlxs

## References

Allen, C.D., Macalady, A.K., Chenchouni, H., Bachelet, D., McDowell, N., Vennetier, M., Kitzberger, T., Rigling, A., Breshears, D.D., Hogg, E.H. (Ted), Gonzalez, P., Fensham, R., Zhang, Z., Castro, J., Demidova, N., Lim, J.H., Allard, G., Running, S.W., Semerci, A. & Cobb, N. (2010) A global overview of drought and heat-induced tree mortality reveals emerging climate change risks for forests. Forest Ecology and Management, 259, 660–684.

Allen, M.R., O.P. Dube, W. Solecki, F. Aragón-Durand, W. Cramer, S. Humphreys, M. Kainuma, J. Kala, N. Mahowald, Y. Mulugetta, R. Perez, M. Wairiu, and K. Zickfeld (2018) Framing and Context. In: Global Warming of 1.5°C. An IPCC Special Report on the impacts of global warming of 1.5°C above pre-industrial levels and related global greenhouse gas emission pathways, in the context of strengthening the global response to the threat of climate change, sustainable development, and efforts to eradicate poverty [Masson-Delmotte, V., P. Zhai, H.-O. Pörtner, D. Roberts, J. Skea, P.R. Shukla, A. Pirani, W. Moufouma-Okia, C. Péan, R. Pidcock, S. Connors, J.B.R. Matthews, Y. Chen, X. Zhou, M.I. Gomis, E. Lonnoy, T. Maycock, M. Tignor, and T. Waterfield (eds.)]. Cambridge University Press, Cambridge, UK and New York, NY, USA, pp. 49–92. 10.1017/9781009157940.003.

Antonelli, A., Fry, C., Smith, R.J., Eden, J., Govaerts, R.H.A., Kersey, P., Nic Lughadha, E., Onstein, R.E., Simmonds, M.S.J., Zizka, A., Ackerman, J.D., Adams, V.M., Ainsworth, A.M., Albouy, C., Allen, A.P., Allen, S.P., Allio, R., Auld. T.D., Bachman, S.P., Baker, W.J., Barrett, R.L., Beaulieu, J.M., Bellot, S., Black, N., Boehnisch, G., Bogarín, D., Boyko, J.D., Brown, M.J.M., Budden, A., Bureš, P., Butt, N., Cabral, A., Cai, L., Cano, J.A., Chang, Y., Charitonidou, M., Chau, J.H., Cheek, M., Chomicki, G., Coiro, M., Colli-Silva, M., Condamine, F.L., Crayn, D.M., Cribb, P., Cuervo-Robayo, A.P., Dahlberg, A., Deklerck, V., Denelle, P., Dhanjal-Adams, K.L., Druzhinina, I., Eiserhardt, W.L., Elliott, T.L., Enquist, B.J., Escudero, M., Espinosa-Ruiz, S., Fay, M.F., Fernández, M., Flanagan, N.S., Forest, F., Fowler, R.M., Freiberg, M., Gallagher, R.V., Gaya, E., Gehrke, B., Gelwick, K., Grace, O.M., Granados Mendoza, C., Grenié, M., Groom, Q.J., Hackel, J., Hagen, E.R., Hágsater, E., Halley, J.M., Hu, A.-Q., Jaramillo, C., Kattge, J., Keith, D.A., Kirk, P., Kissling, W.D., Knapp, S., Kreft, H., Kuhnhäuser, B.G., Larridon, I., Leão, T.C.C., Leitch, I.J., Liimatainen, K., Lim, J.Y., Lucas, E., Lücking, R., Luján, M., Luo, A., Magallón, S., Maitner, B., Márquez-Corro, J.I., Martín-Bravo, S., Martins-Cunha, K., Mashau, A.C., Mauad, A.V., Maurin, O., Medina Lemos, R., Merow, C., Michelangeli, F.A., Mifsud, J.C.O., Mikryukov, V., Moat, J., Monro, A.K., Muasya, A.M., Mueller, G.M., Muellner-Riehl, A.N., Nargar, K., Negrão, R., Nicolson, N., Niskanen, T., Oliveira Andrino, C., Olmstead, R.G., Ondo, I., Oses, L., Parra-Sánchez, E., Paton, A.J., Pellicer, J., Pellissier, L., Pennington, T.D., Pérez-Escobar, O.A., Phillips, C., Pironon, S., Possingham, H., Prance, G., Przelomska, N.A.S., Ramírez-Barahona, S.A., Renner, S.S., Rincon, M., Rivers, M.C., Rojas Andrés, B.M., RomeroSoler, K.J., Roque, N., Rzedowski, J., Sanmartín, I., Santamaría-Aguilar, D., Schellenberger Costa, D., Serpell, E., Seyfullah, L.J., Shah, T., Shen, X., Silvestro, D., Simpson, D.A., Šmarda, P., Šmerda, J., Smidt, E., Smith, S.A., Solano-Gomez, R., Sothers, C., Soto Gomez, M., Spalink, D., Sperotto, P., Sun, M., Suz, L.M., Svenning, J.-C., Taylor, A., Tedersoo, L., Tietje, M., Trekels, M., Tremblay, R.L., Turner, R., Vasconcelos, T., Veselý, P., Villanueva, B.S., Villaverde, T., Vorontsova, M.S., Walker, B.E., Wang, Z., Watson, M., Weigelt, P., Wenk, E.H., Westrip, J.R.S., Wilkinson, T., Willett, S.D., Wilson, K.L., Winter, M., Wirth, C., Wölke, F.J.R., Wright, I.J., Zedek, F., Zhigila, D.A., Zimmermann, N.E., Zuluaga, A., Zuntini, A.R. (2023). State of the World’s Plants and Fungi 2023. Royal Botanic Gardens, Kew. DOI: 10.34885/wnwn-6s63

Araújo, M.B., Ferri-Yáñez, F., Bozinovic, F., Marquet, P. a, Valladares, F. & Chown, S.L. (2013) Heat freezes niche evolution. Ecology Letters, 16, 1206–19.

Baker, N.R. (2008) Chlorophyll Fluorescence: A Probe of Photosynthesis In Vivo. Annual Review of Plant Biology, 59, 89–113.

Bates, D., Mächler, M., Bolker, B., & Walker, S. (2015). Fitting linear mixed-effects models using lme4. Journal of Statistical Software, 67(1), 1–48. 10.18637/jss.v067.i01

Bennett, J.M., Calosi, P., Clusella-Trullas, S., Martínez, B., Sunday, J., Algar, A.C., Araújo, M.B., Hawkins, B.A., Keith, S., Kühn, I., Rahbek, C., Rodríguez, L., Singer, A., Villalobos, F., Ángel Olalla-Tárraga, M. & Morales-Castilla, I. (2018) GlobTherm, a global database on thermal tolerances for aquatic and terrestrial organisms. Scientific Data, 5, 1–7.

Biebl, R. (1964) Temperaturresistenz tropischer Pflanzen auf Puerto Rico. Protoplasma, 59, 133–156.

Biebl, R. & Maier, R. (1969) Tageslänge und Temperaturresistenz. Österr. Bot. Z., 117, 176– 194.

Bison, N.N. & Michaletz, S.T. (2024) Variation in leaf carbon economics, energy balance, and heat tolerance traits highlights differing timescales of adaptation and acclimation. New Phytologist, 242, 1919–1931.

Bita, C.E. & Gerats, T. (2013) Plant tolerance to high temperature in a changing environment: scientific fundamentals and production of heat stress-tolerant crops. Frontiers in Plant Science, 4, 1–18.

Brown, M.J.M., Walker, B.E., Black, N., Govaerts, R.H.A., Ondo, I., Turner, R. & Nic Lughadha, E. (2023) R WCVP: a companion R package for the World Checklist of Vascular Plants. New Phytologist, 240, 1355–1365.

Buchner, O. & Neuner, G. (2003) Variability of Heat Tolerance in Alpine Plant Species Measured at Different Altitudes. Arctic, Antarctic, and Alpine Research, 35, 411–420.

Buchner, O., Roach, T., Gertzen, J., Schenk, S., Karadar, M., Stöggl, W., Miller, R., Bertel, C., Neuner, G. & Kranner, I. (2017) Drought affects the heat-hardening capacity of alpine plants as indicated by changes in xanthophyll cycle pigments, singlet oxygen scavenging, α-tocopherol and plant hormones. Environmental and Experimental Botany, 133, 159– 175.

Chen, H.-H., Shen, Z.-Y. & Li, P.H. (1982) Adaptability of Crop Plants to High Temperatures Stress. Crop Science, 22, 719.

Colombo, S.J. & Timmer, V.R. (1992) Limits of tolerance to high temperatures causing direct and indirect damage to white spruce. Tree Physiology, 11, 95–104.

Cook, A.M., Rezende, E.L., Petrou, K. & Leigh, A. (2024) Beyond a single temperature threshold: Applying a cumulative thermal stress framework to plant heat tolerance. Ecology Letters, 27, e14416.

Crop Wild Relatives Occurrence data consortia (2024). A global database for the distributions of crop wild relatives. Version 1.13. Centro Internacional de Agricultura Tropical - CIAT. Occurrence dataset 10.15468/jyrthk accessed via GBIF.org on 2025-08-31.

Doughty, C.E., Keany, J.M., Wiebe, B.C., Rey-Sanchez, C., Carter, K.R., Middleby, K.B., Cheesman, A.W., Goulden, M.L., da Rocha, H.R., Miller, S.D., Malhi, Y., Fauset, S., Gloor, E., Slot, M., Oliveras Menor, I., Crous, K.Y., Goldsmith, G.R. & Fisher, J.B. (2023) Tropical forests are approaching critical temperature thresholds. Nature, 621, 105–111.

Endris, J. & Rehm, E. (2024) Leaf temperatures exceed thermal heat tolerances for a community of eastern North America hardwood trees. AoB PLANTS, plae060.

Feeley, K., Martinez-villa, J., Perez, T. & Duque, A.S. (2020) The thermal tolerances,distributions, and performances of tropical montane tree species. Frontiers in Forests and Global Change, 3, 1–11.

Gauslaa, Y. (1984) Heat Resistance and Energy Budget in Different Scandinavian Plants. Holarctic Ecology, 7, 5–78.

Geange, S.R., Arnold, P.A., Catling, A.A., Coast, O., Cook, A.M., Gowland, K.M., Leigh, A., Notarnicola, R.F., Posch, B.C., Venn, S.E., Zhu, L. & Nicotra, A.B. (2021) The thermal tolerance of photosynthetic tissues: a global systematic review and agenda for future research. New Phytologist, 229, 2497–2513.

Godoy, O., Lemos-filho, J.P.D. & Valladares, F. (2011) Invasive species can handle higher leaf temperature under water stress than Mediterranean natives Author’ s personal copy. Environmental and Experimental Botany, 71, 207–214.

Govaerts, R., Nic Lughadha, E. et al. The World Checklist of Vascular Plants, a continuously updated resource for exploring global plant diversity. Sci Data 8, *215* (2021).

Hauck, M., Schneider, T., Bahlinger, S., Fischbach, J., Oswald, G., Csapek, G. & Dulamsuren, C. (2025) Heat tolerance of temperate tree species from Central Europe. Forest Ecology and Management, 580, 122541.

Havaux, M. (1993) Rapid photosynthetic adaptation to heat stress triggered in potato leaves by moderately elevated temperatures. *Plant*, Cell & Environment, 16, 461–467.

Hernández, G.G., Perez, T.M., Vargas, O.M., Kress, W.J., Molina Bravo, R., Cordero, R.A., Seemann, J.R. & García Robledo, C. (2022) Evolutionary history constrains heat tolerance of native and exotic tropical Zingiberales. Functional Ecology, 36, 3073–3084.

Hüve, K., Bichele, I., Rasulov, B. & Niinemets, Ü. (2011) When it is too hot for photosynthesis: heat induced instability of photosynthesis in relation to respiratory burst, cell permeability changes and H_2_ O_2_ formation. *Plant*, Cell & Environment, 34, 113–126.

Iturbide, M., Gutiérrez, J.M., Alves, L.M., Bedia, J., Cerezo-Mota, R., Cimadevilla, E., Cofiño, A.S., Di Luca, A., Faria, S.H., Gorodetskaya, I.V., Hauser, M., Herrera, S., Hennessy, K., Hewitt, H.T., Jones, R.G., Krakovska, S., Manzanas, R., Martínez-Castro, D., Narisma, G.T., Nurhati, I.S., Pinto, I., Seneviratne, S.I., Van Den Hurk, B. & Vera, C.S. (2020) An update of IPCC climate reference regions for subcontinental analysis of climate model data: definition and aggregated datasets. Earth System Science Data, 12, 2959–2970.

Kitao, M., Lei, T.T., Koike, T., Matsumoto, Y., Ang, L.-H., Tobita, H. & Maruyama, Y. (2000) Temperature response and photoinhibition investigated by chlorophyll fluorescence measurements for four distinct species of dipterocarp trees. Physiologia Plantarum, 109, 284–290.

Knight, C.A. & Ackerly, D.D. (2003) Evolution and plasticity of photosynthetic thermal tolerance, specific leaf area and leaf size: Congeneric species from desert and coastal environments. New Phytologist, 160, 337–347.

Krause, G.H. & Santarius, K.A. (1975) Relative Thermostability of the Chloroplast Envelope. Planta, 299.

Krause, G.H., Winter, K., Krause, B., Jahns, P., García, M., Aranda, J. & Virgo, A. (2010) High-temperature tolerance of a tropical tree, Ficus insipida: methodological reassessment and climate change considerations. Functional Plant Biology, 37, 890.

Krause, G.H., Winter, K., Krause, B. & Virgo, A. (2015) Light-stimulated heat tolerance in leaves of two neotropical tree species, Ficus insipida and Calophyllum longifolium. Functional Plant Biology, 42, 42–51.

Kullberg, A.T. & Feeley, K.J. (2024) Seasonal acclimation of photosynthetic thermal tolerances in six woody tropical species along a thermal gradient. Functional Ecology, 2493–2505.

Lancaster, L.T. & Humphreys, A.M. (2020) Global variation in the thermal tolerances of plants. Proceedings of the National Academy of Sciences, 1–8.

Lange, O.L. (1965) *THE HEAT RESISTANCE OF PLANTS, ITS DETERMINATION AND VARIABILITY*. *METHODOLOGY OF PLANT ECO-PHYSIOLOGY*, pp. 399–405. United nations Educational, Scientific and Cultural Organization.

Langridge, P. & Reynolds, M. (2021) Breeding for drought and heat tolerance in wheat. Theoretical and Applied Genetics, 134, 1753–1769.

Larcher, W., Holzner, M. & Pichler, J. (1989) Temperaturresistenz inneralpiner Trockenrasen. Flora, 183, 115–131.

Leigh, A., Sevanto, S., Ball, M.C., Close, J.D., Ellsworth, D.S., Knight, C.A., Nicotra, A.B. & Vogel, S. (2012) Do thick leaves avoid thermal damage in critically low wind speeds? New Phytologist, 194, 477–487.

Lutterschmidt, W.I. & Hutchison, V.H. (1997) The critical thermal maximum: History and critique. Canadian Journal of Zoology, 75, 1561–1574.

Mooney, H.A. & Billings, W.D. (1961) Comparative Physiological Ecology of Arctic and Alpine Populations of Oxyria digyna. Ecological Monographs, 31, 1–29.

Neri, P., Gu, L. & Song, Y. (2024) The effect of temperature on photosystem II efficiency across plant functional types and climate. Biogeosciences, 21, 2731–2758.

Nobel, P.S. (1984) Oecologia Extreme temperatures and thermal tolerances for seedlings of desert succulents. 310–317.

Offord, C. a. (2011) Pushed to the limit: Consequences of climate change for the Araucariaceae: A relictual rain forest family. Annals of Botany, 108, 347–357.

Onwueme, I.C. (1979) Rapid, plant-conserving estimation of heat tolerance in plants. Journal of Agricultural Sciences, 92, 527–535.

O’Sullivan, O.S., Heskel, M.A., Reich, P.B., Tjoelker, M.G., Weerasinghe, K.W.L.K., Penillard, A., Zhu, L., Egerton, J.J.G., Bloomfield, K.J., Creek, D., Bahar, N.H.A., Griffin, K.L., Hurry, V., Meir, P., Turnbull, M.H. & Atkin, O.K. (2017) Thermal limits of leaf metabolism across biomes. Global Change Biology, 209–223.

Perez, T. M. (2019). The ecophysiology of photosynthetic heat tolerances in tropical plants [Doctoral dissertation]. University of Miami.

Perez, T. & Feeley, K. (2020) Photosynthetic heat tolerances and extreme leaf temperatures. Functional Ecology, 34, 2236–2245.

Perez, T.M. (2020) Weak phylogenetic and climatic signals in plant heat tolerance. Journal of Biogeography, 1–10.

Perez, T.M., Feeley, K.J., Michaletz, S.T. & Slot, M. (2021) Methods matter for assessing global variation in plant thermal tolerance. Proceedings of the National Academy of Sciences of the United States of America, 118, 10–11.

Rehfeldt, G.E., Leites, L.P., Bradley St Clair, J., Jaquish, B.C., Sáenz-Romero, C., López-Upton, J. & Joyce, D.G. (2014) Comparative genetic responses to climate in the varieties of Pinus ponderosa and Pseudotsuga menziesii: Clines in growth potential. Forest Ecology and Management, 324, 138–146.

Sapper, I. (1935) Versuche zur hitzeresistenz der pflanzen. Planta, 23, 518–556.

Schreiber, U., Colbow, K. & Vidaver, W. (1976) Analysis of temperature-jump chlorophyll fluorescence induction in plants. Biochimica et Biophysica Acta, 423, 249–263.

Slot, M., Cala, D., Aranda, J., Virgo, A., Michaletz, S.T. & Winter, K. (2021) Leaf heat tolerance of 147 tropical forest species varies with elevation and leaf functional traits, but not with phylogeny. Plant, Cell & Environment, 44, 2414–2427.

Valentin tefan, & Sam Levin. (2018). plotbiomes: R package for plotting Whittaker biomes with ggplot2 (v1.0.0). Zenodo. 10.5281/zenodo.7145245

Tarvainen, L., Wittemann, M., Mujawamariya, M., Manishimwe, A., Zibera, E., Ntirugulirwa, B., Ract, C., Manzi, O.J.L., Andersson, M.X., Spetea, C., Nsabimana, D., Wallin, G. & Uddling, J. (2022) Handling the heat – photosynthetic thermal stress in tropical trees. New Phytologist, 233, 236–250.

Wahid, A., Gelani, S., Ashraf, M. & Foolad, M. (2007) Heat tolerance in plants: An overview. Environmental and Experimental Botany, 61, 199–223.

Winter, K., Krüger Nuñez, C.R., Slot, M. & Virgo, A. (2024) In thermotolerance tests of tropical tree leaves, the chlorophyll fluorescence parameter Fv/Fm measured soon after heat exposure is not a reliable predictor of tissue necrosis. Plant Biology, 27, 146–153.

Zhu, L., Bloomfield, K.J., Hocart, C.H., Egerton, J.J.G., O’Sullivan, O.S., Penillard, A., Weerasinghe, L.K. & Atkin, O.K. (2018) Plasticity of photosynthetic heat tolerance in plants adapted to thermally contrasting biomes. Plant Cell and Environment, 41, 1251– 1262.

Zhu, L., Scafaro, A.P., Vierling, E., Ball, M.C., Posch, B.C., Stock, F. & Atkin, O.K. (2024) Heat tolerance of a tropical–subtropical rainforest tree species Polyscias elegans: time dependent dynamic responses of physiological thermostability and biochemistry. New Phytologist, 241, 715–731.

