## Supplemental c for "A database of plant heat tolerances and methodological matters"

Supporting Information:

**Section 1: Literature Search:**

The Perez 2019 dataset was cross-referenced against the Lancaster and Humphreys (2020) dataset, and incorporated into the first version of our dataset. Additional references were obtained through forward and backward references searches and their data was incorporated our dataset. The resulting dataset labeled Perez et al 2025 V1was submitted for review.

Following the first review, we cross-referenced the GlobTherm dataset and incorporated data from applicable references into our database. An additional 3 references from the Lancaster and Humphreys dataset were also incorporated at this time. The resulting database in termed Perez et al 2025 V2.

Here, we describe how we incorporated data from references and how our collection procedures led to notable deviations in the data we reported compared to other datasets. For example, the GlobTherm dataset contained 12 references corresponding to 73 heat tolerance records for 73 species after filtering for “Tmax” and “Streptophytes” (Table 1). Seven of these references were already common to the Perez 2019 dataset. These seven references corresponded to 222 heat tolerance records for 81, and 63 records from 62 species in the Perez 2019 and GlobTherm datasets, respectively (main text Table 2). We incorporated data from 4 new references unique to the Globtherm dataset into our current dataset, which corresponded to 14 heat tolerances records for 11 species. One reference in the GlobTherm database was excluded because it provided data for algae and our study focuses on terrestrial plants.

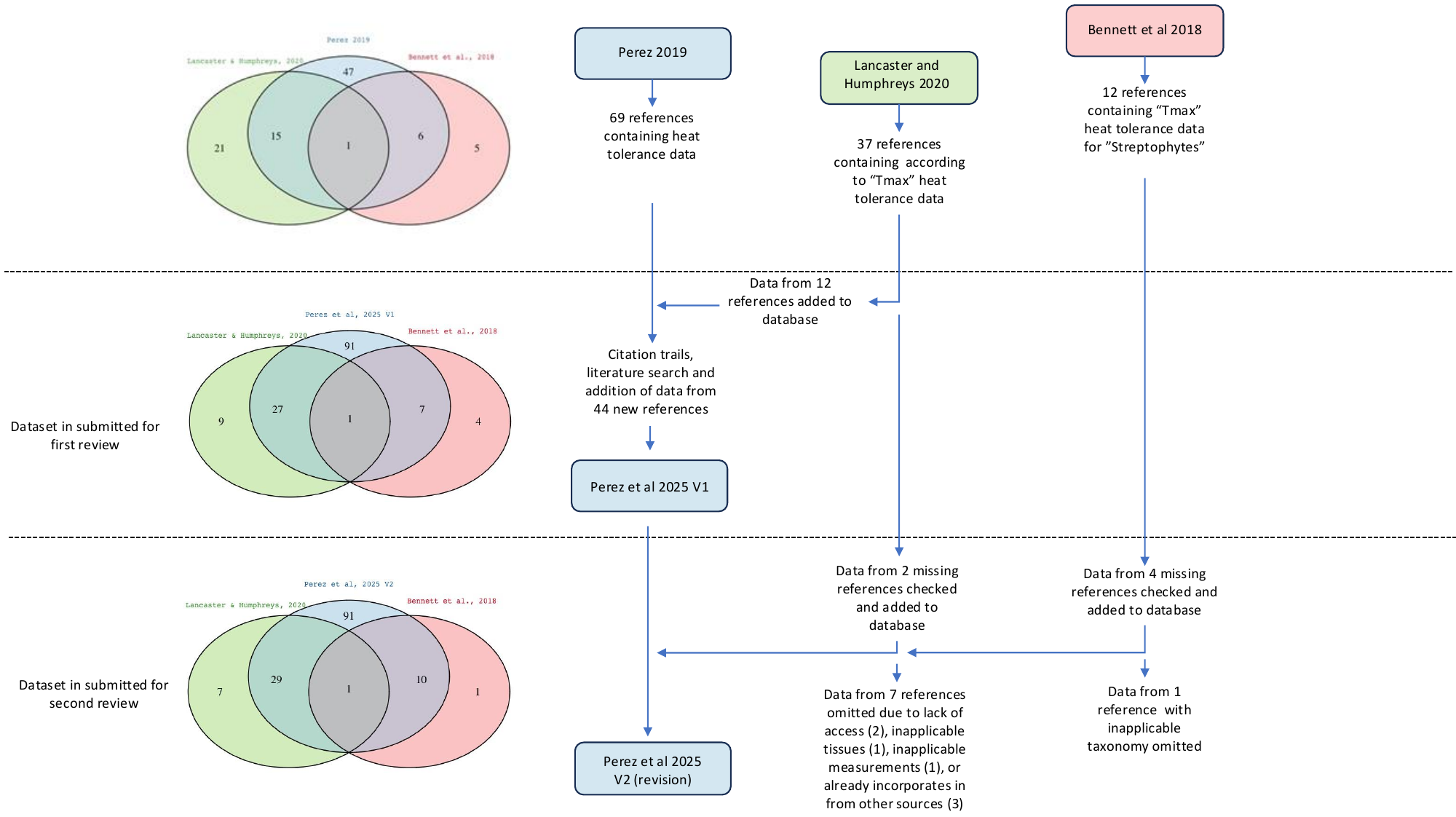

Figure S1: A diagram depicting how references from Perez 2019, Lancaster and Humphrey (2020), and the GlobTherm (Bennett 2017) databases were integrated.

We filtered the Lancaster and Humphreys (2020) dataset to include data categorized as “Tmax”, which yielded 37 applicable references (a total of 689 species and 964 records; Table 1), of which 16 were common to the Perez 2019 dataset. The 16 overlapping references represented 716 records for 566 species and 778 records for 545 species for the Lancaster and Humphreys (2020) and Perez 2019, datasets respectively. We then incorporated data from 14 references unique to the Lancaster and Humphreys (2020) into our database, which corresponded to 128 new heat tolerance records for 77 species in our database (Table 2). We disqualified 7 references included in the Lancaster and Humphreys dataset because they had 1) data already incorporated into our database but sourced from other references, 2) did not have analogous physiological measurements (e.g. carbon assimilation or respiration data), 3) were taxonomically ineligible algae, or 4) could not be obtained for assessment.

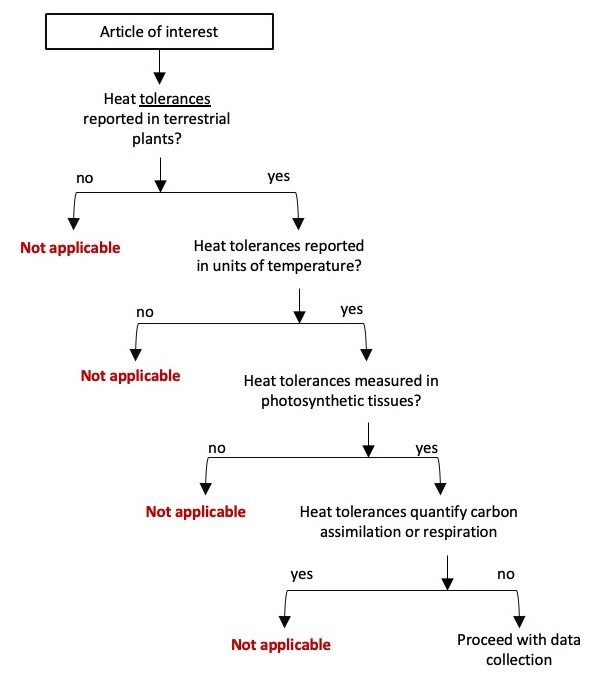

*Figure S2:* Diagram illustrating the criteria we used to collect data from articles

We followed a decision tree, illustrated in Figure S2, to determine if data were eligible to be included in database. This process generally involved reading the articles to clearly understand the data different studies were reporting, and in doing so provided references to other studies that we collected data from. Studies that we deemed eligible were reviewed more thoroughly for full data collection efforts that are described in the Methods sections of our main text.

***Section 2: Methodological data descriptions, results and discussion***

*Table S3:* The categorized standardized terminology and the original terminology reported within our database. Superscripts correspond to references provided below.

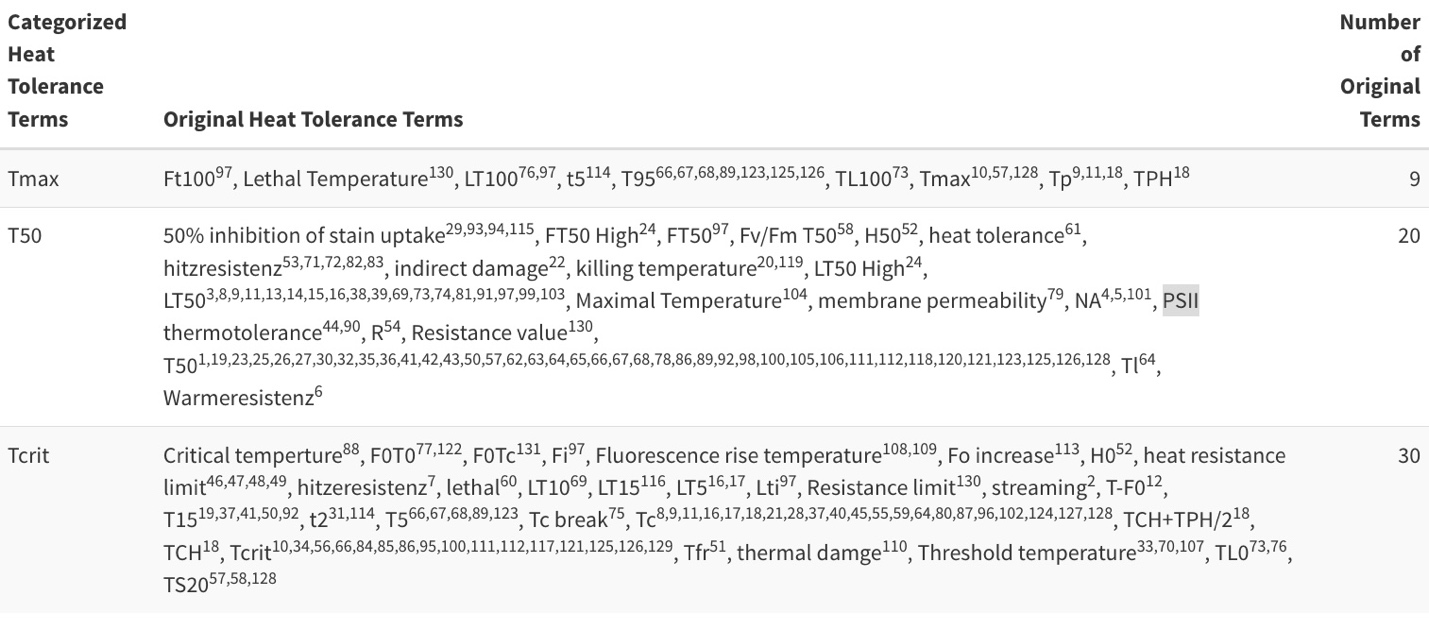

**Reference list:**

1: Ahrens CW, Challis A, Byrne M, Leigh A, Nicotra AB, Tissue D, Rymer P. 2021. Repeated extreme heatwaves result in higher leaf thermal tolerances and greater safety margins. New Phytologist 232: 1212-1225

2: Alexandrov VY. 1964. Cytophysiological and Cytoecological Investigations of Heat Resistance of Plant Cells Toward the Action of High and Low Temperature. The Quarterly Review of Biology 39: 35-77

3: Balagurova, N., Drozdov, S. & Grabovik, S. 1996. Cold and heat resistance of five species of Sphagnum. Annales Botanici Fennici. 33, 33-37.

4: Berry, J, Bjorkman, O. 1980. Photosynthetic Response and Adaptation to Temperature in Higher Plants. Annu. Rev. Plant Physiol. 31: 491-543

5: Biebl R, Maier. 1969. Tageslange und Temperaturresistenz. Osterreichische Botanische Zeitschrift, 117(2), 176-194

6: Biebl, R. 1964. Temperaturresistenz tropischer Pflanzen auf Puerto Rico. Protoplasma 59: 133-156

7: Biebl, R. 1967. √úber W√§rmehaushalt und Temperaturresistenz arktischer Pflanzen in Westgr√∂nland. Flora oder Allg. Bot. Zeitung. Abt. B, Morphol. und Geobot. 157: 327‚Äì354. Urban & Fischer.

8: Bigras FJ. 2000. Selection of white spruce families in the context of climate change: Heat tolerance. Tree Physiology 20: 1227-1234

9: Bilger H-W, Schreiber U, Lange OL. 1984. Determination of leaf heat resistance: comparative and tissue necrosis methods. Oecologia 63: 256-262

10: Bison, N, Michaletz, S. 2024. Variation in leaf carbon economics, energy balance, and heat tolerance traits highlights differing timescales of adaptation and acclimation. New Phytologist https://doi.org/10.1111/nph.19702

11: Braun, V, Buchner, O, Neuner, G. 2002. Thermotolerance of Photosystem 2 of Three Alpine Plant Species Under Field Conditions. Photosynthetica 40: 587-595. Kluwer Academic Publishers.

12: Bryant C, Harris RJ, Brothers N, Bone C, Walsh N, Nicotra AB, Ball MC. 2024. Cross-tolerance: Salinity gradients and dehydration increase photosynthetic heat tolerance in mangrove leaves. Functional Ecology 00: 1-13

13: Buchner O, Karadar M, Bauer I, Neuner G. 2013. A novel system for in situ determination of heat tolerance of plants: first results on alpine dwarf shrubs. Plant methods 9: 7

14: Buchner O, Neuner G. 2001. Determination of heat tolerance: A new equipment for field measurements. Journal of Applied Botany and Food Quality 75: 130-137

15: Buchner O, Neuner G. 2003. Variability of Heat Tolerance in Alpine Plant Species Measured at Different Altitudes. Arctic, Antarctic, and Alpine Research 35: 411-420

16: Buchner, O, Roach, T, Gertzen, J, Schenk, S, Karadar, M, St√∂ggl, W, Miller, R, Bertel, C, Neuner, G, Kranner, I. 2017. Drought affects the heat-hardening capacity of alpine plants as indicated by changes in xanthophyll cycle pigments, singlet oxygen scavenging, a-tocopherol and plant hormones. Environmental and Experimental Botany. 133: 159-177

17: Buchner, O, Stoll, M, Karadar, M, Kranner, I, Neuner, G. 2015. Application of heat stress in situ demonstrates a protective role of irradiation on photosynthetic performance in alpine plants. Plant. Cell Environ. 38: 812-26

18: Buchner, O. & Neuner, G. 2010. Freezing cytorrhysis and critical temperature thresholds for photosystem II in the peat moss Sphagnum capillifolium. Protoplasma 243, 63-71.

19: Chaves, CJN, Leal, BSS, Lemos-Filho, JP de. 2018. How are endemic and widely distributed bromeliads responding to warming temperatures? A case study in a Brazilian hotspot. Flora 238: 110-118

20: Chen H-H, Shen Z-Y, Li PH. 1982. Adaptability of Crop Plants to High Temperatures Stress. Crop Science 22: 719

21: Choinski Jr, JS, Gould, KS. 2010. Immature leaves of Weinmannia racemosa are more heat tolerant than mature leaves based on differences in chlorophyll a fluorescence and solute leakage. New Zealand Journal of Botany. 48(3-4): 163-177

22: Colombo SJ, Timmer VR. 1992. Limits of tolerance to high temperatures causing direct and indirect damage to white spruce. Tree Physiology 11: 95-104

23: Cook, AM, Berry, N, Milner, KV, Leigh, A. 2021. Water availability influences thermal safety margins for leaves. Functional Ecology. 35: 2179-2189

24: Cunningham SC, Read J. 2006. Foliar temperature tolerance of temperate and tropical evergreen rain forest trees of Australia. Tree Physiology 26: 1435-1443

25: Curtis EM, Gollan J, Murray BR, Leigh A. 2016. Native microhabitats better predict tolerance to warming than latitudinal macro-climatic variables in arid-zone plants. Journal of Biogeography 43: 1156-1165

26: Curtis EM, Knight CA, Leigh A. 2019. Intracanopy adjustment of leaf-level thermal tolerance is associated with microclimatic variation across the canopy of a desert tree (Acacia papyrocarpa). Oecologia 189: 37-46

27: Curtis EM, Knight CA, Petrou K, Leigh A. 2014. A comparative analysis of photosynthetic recovery from thermal stress: A desert plant case study. Oecologia 175: 1051-1061

28: Daas C, Montpied P, Hanchi B, Dreyer E. 2008. Responses of photosynthesis to high temperatures in oak saplings assessed by chlorophyll-a fluorescence: inter-specific diversity and temperature-induced plasticity. Annals of Forest Science 65: 305

29: Didden-zopfy, B, & Nobel, P S. 1982. Opuntia bigelovii. Oecologia, 176-180

30: Docherty EM, Gloor E, Sponchiado D, Gilpin M, Pinto CAD, Junior HM, Coughlin I, Ferreira L, Junior JAS, da Costa ACL, Meir P, Galbraith D. 2022. Long-term drought effects on the thermal sensitivity of Amazon forest trees. Plant, Cell & Environment 46(1):185-198

31: Downton WJS, Berry JA, Seemann JR. 1984. Tolerance of Photosynthesis to High Temperature in Desert Plants. Plant Physiology 74: 786-790

32: Drake JE, Tjoelker MG, V?rhammar A, Medlyn BE, Reich PB, Leigh A, Pfautsch S, Blackman CJ, Lopez R, Aspinwall MJ, et al. 2018. Trees tolerate an extreme heatwave via sustained transpirational cooling and increased leaf thermal tolerance. Global Change Biology 24: 2390-2402

33: Epron D. 1997. Effects of drought on photosynthesis and on the thermotolerance of photosystem II in seedlings of cedar (Cedrus atlantica and C. libani ). Journal of Experimental Botany 48: 1835-1841

34: Esperon-Rodriguez, M, Power, SA, Tjoelker, MG, Marchin, RM, Rymer, PD. 2021. Contrasting heat tolerance of urban trees to extreme temperatures during heatwaves. Urban Forestry and Urban Greeing. 66:127387

35: Fadrique, B, Baraloto, C, Bravo-Avila, C, Feeley, K. 2022. Bamboo climatic tolerances are decoupled from leaf functional traits across an Andean elevation gradient. Oikos. e09229

36: Feeley, K, Martinez-villa, J, Perez, T, & Duque, AS. 2020. The thermal tolerances,distributions ,and performances of tropical montane tree species. Frontiers in Forests and Global Change 3: 1-11. https://doi.org/10.3389/ffgc.2020.00025

37: Froux F, Ducrey M, Epron D, Dreyer E, Froux F, Ducrey M, Epron D, Dreyer E, Rouxa FF, Ucreyb MD, et al. 2004. Seasonal variations and acclimation potential of the thermostability of photochemistry in four Mediterranean conifers. Ann. For. Sci.: 235-241

38: Garrett, TY, Huynh, C-V, North, GB. 2010. Root contraction helps protect the ‚Äôliving rock‚Äú cactus Ariocarpus fissuratus from lethal high temperatures when growing in rocky soil. Am. J. Bot. 97: 1951-1960

39: Gauslaa Y. 1984. Heat Resistance and Energy Budget in Different Scandinavian Plants. Holarctic Ecology 7: 5-78

40: Ghouil H, Montpied P, Epron D, Ksontini M, Hanchi B, Dreyer E. 2003. Thermal optima of photosynthetic functions and thermostability of photochemistry in cork oak seedlings. Tree Physiology: 1031-1039

41: Gimeno TE, Pias B, Lemos-Filho J, Valladares F. 2008. Plasticity and stress tolerance override local adaptation in the responses of Mediterranean holm oak seedlings to drought and cold. Tree Physiology: 87-98

42: Godoy O, Lemos-filho JP De, Valladares F. 2011. Invasive species can handle higher leaf temperature under water stress than Mediterranean natives. Environmental and Experimental Botany 71: 207-214

43: Hanley, PA, Arndt, SK, Livesley, SJ, Szota, C. 2021. Relating the climate envelopes of urban tree species to their drought and thermal tolerance. Science of the total environment. 753: 142012

44: Havaux M, Tardy F. 1996. Temperature-dependent adjustment of the thermal stability of photosystem II in vivo: possible involvement of xanthophyll-cycle pigments. Planta 198: 324-333

45: Havaux M. 1993. Rapid photosynthetic adaptation to heat stress triggered in potato leaves by moderately elevated temperatures. Plant, Cell & Environment 16: 461-467

46: Hellmuth EO. 1968. Eco-Physiological Studies on Plants in Arid and Semi-Arid Regions in Western Australia: I. Autecology of Rhagodia Baccata (Labill.) Moq. The Journal of Ecology 56: 319-344

47: Hellmuth EO. 1971 a. Eco-Physiological Studies on Plants in Arid and Semi-Arid Regions in Western Australia: V. Heat Resistance Limits of Photosynthetic Organs of Different Seasons, Their Relation to Water Deficits and Cell Sap Properties and the Regeneration Ability. The Journal of Ecology 59: 365-374

48: Hellmuth EO. 1971 b. Eco-Physiological Studies on Plants in Arid and Semi-Arid Regions in Western Australia: IV. Comparison of the Field Physiology of the Host, Acacia Grasbyi and its Hemiparasite , Amyema Nestor Under Optimal and Stress Conditions. Journal of Ecology 59: 351-363

49: Hellmuth EO. 1971 c. Eco-Physiological Studies on Plants in Arid and Semi-Arid Regions in Western Australia: III. Comparative Studies on Photosynthesis, Respiration and Water Relations of Ten Arid Zone and Two Semi-Arid Zone Plants Under Winter and Late Summer Climatic Condit. Journal of Ecology 59: 225-259

50: Hernandez, G, Perez, T, Vargas, O, Kress, W, Molina-Bravo, R, Cordero, R, Seemann, J, Garcia-Robledo, C. 2022 Evolutionary history constrains heat tolerance of native and exotic tropical Zingiberales. Functional Ecology 36:3073‚Äì3084

51: Huve K, Bichele I, Tobias M, Niinemets. 2006. Heat sensitivity of photosynthetic electron transport varies during the day due to changes in sugars and osmotic potential. Plant, Cell and Environment 29: 212-228

52: Ishikawa, M, Gusta, LV. 1996. Freezing and heat tolerance of Opuntia cacti native to the Canadian prairie provinces. Canadian Journal of Botany. 74:1890-1895

53: Kappen L. 1964. Untersuchungen fiber den Jahreslauf. der Frost-, Hitze- und Austrocknungsresistenz von Sporophyten einheimischer Polypodiaceen (Filicinae) Flora, 155, 123-166

54: Karschon R, Pinchas L. 1971. Variations In Heat Resistance of Ecotypes. Aust. J. Bot.: 261-272

55: Kitao M, Lei TT, Koike T, Matsumoto Y, Ang L-H, Tobita H, Maruyama Y. 2000. Temperature response and photoinhibition investigated by chlorophyll fluorescence measurements for four distinct species of dipterocarp trees. Physiologia Plantarum 109: 284-290

56: Kitudom, N, Fauset, S, Zhou, Y, Fan, Z, Li, M, He, M, Zhang, S, Xu, K, Lin, H. 2022. Thermal safety margins of plant leaves across biomes under a heatwave. Science of the Total Environment 806:150416

57: Knight CA, Ackerly DD. 2002. An ecological and evolutionary analysis of photosynthetic thermotolerance using the temperature-dependent increase in fluorescence. Oecologia 130: 505-514

58: Knight CA, Ackerly DD. 2003. Evolution and plasticity of photosynthetic thermal tolerance, specific leaf area and leaf size: Congeneric species from desert and coastal environments. New Phytologist 160: 337-347

59: Koniger, M, Harris, GC, Pearcy, RW. 1998. Interaction between photon flux density and elevated temperatures on photoinhibition in Alocasia macrorrhiza. Planta. 205:214-222

60: Konis, E. 1949. The Resistance of Maquis Plants to Supramaximal Temperatures. Ecology, 30: 425-429. https://doi.org/10.2307/1932445

61: Krause GH, Cheesman AW, Winter K, Krause B, Virgo A. 2013. Thermal tolerance, net CO2 exchange and growth of a tropical tree species, Ficus insipida, cultivated at elevated daytime and nighttime temperatures. Journal of Plant Physiology 170: 822-827

62: Krause GH, Winter K, Krause B, Virgo A. 2015. Light-stimulated heat tolerance in leaves of two neotropical tree species , Ficus insipida and Calophyllum longifolium. Functional Plant Biology 42: 42-51

63: Krause GH, Winter K, Krause B, Virgo A. 2016. Protection by light against heat stress in leaves of tropical crassulacean acid metabolism plants containing high acid levels. Functional Plant Biology 43: 1061-1069

64: Krause, G.H., Winter, K., Krause, B., Jahns, P., Garc‚Äôa, M., Aranda, J., et al. 2010. High-temperature tolerance of a tropical tree, Ficus insipida: Methodological reassessment and climate change considerations. Funct. Plant Biol. 37: 890-900

65: Krause, GH, Cheesman, AW, Winter, K, Krause, B, Virgo, A. 2013. Thermal tolerance, net CO2 exchange and growth of a tropical tree species, Ficus insipida, cultivated at elevated daytime and nighttime temperatures. Journal of Plant Physiology. 170:822-827

66: Kullberg AT, Coombs L, Soria Ahuanari RD, Fortier RP, Feeley KJ. 2023. Leaf thermal safety margins decline at hotter temperatures in a natural warming experiment in the Amazon. New Phytologist. doi: 10.1111/nph.19413

67: Kullberg AT, Feeley KJ. 2022. Limited acclimation of leaf traits and leaf temperatures in a subtropical urban heat island. Tree Physiology 42(11):2266‚Äì2281

68: Kunert, N, Hajek, P, Hietz, P, Morris, H, Rosner, S, Tholen, D. 2022. Summer temperatures reach the thermal tolerance threshold of photosynthetic decline in temperate conifers. Plant Biology. 24:1254-1261

69: Ladinig, U, Pramsohler, M, Bauer, I, Zimmermann, S, Neuner, G, Wagner, J. 2015. Is sexual reproduction of high-mountain plants endangered by heat. Oecologia 177:1195-2010

70: Ladjal M, Epron D, Ducrey M. 2000. Effects of drought preconditioning on thermotolerance of photosystem II and susceptibility of photosynthesis to heat stress in cedar seedlings. Journal of Experimental Botany 20: 1235-1241

71: Lange OL, Lange R. 1963. Untersuchungen fiber Blattemperaturen, Transpiration und Hitzeresistenz an Pflanzen mediterraner Standorte (Costa brava, Spanien). Flora 153: 387-425

72: Lange OL. 1959. Untersuchungen tiber Warmehaushalt und Hitzeresistenz mauretanischer Wtistenund Savannenpflanzen. Flora oder Allgemeine botanische Zeitung, 147: 595-651

73: Larcher W, Holzner M, Pichler J. 1989. Temperaturresistenz inneralpiner Trockenrasen. Flora 183: 115-131

74: Larcher W, Kainmuller C, Wagner J. 2010. Survival types of high mountain plants under extreme temperatures. Flora: Morphology, Distribution, Functional Ecology of Plants 205: 3-18

75: Larcher W, Wagner J, Thammathaworn A. 1990. Effects of Superimposed Temperature Stress on in vivo Chlorophyll Fluorescence of Vigna unguiculata under Saline Stress. Journal of Plant Physiology 136: 92-102

76: Larcher W, Wagner J. 1976. Temperaturgrenzen der CO2-Aufnahme und Temperaturresistenz der Blaetter von Gebirgspflanzen im vegetationsaktiven Zustand. Oecologia Plantarum 11: 361-374

77: Larcher, W, Wagner, J, Lutz C. 1997. The effect of heat on photosynthesis, dark respiration and cellular ultrastructure of the arctic-alpine psychrophyte Ranunculus glacialis 34: 219-232

78: Leon-garcia, IV, & Lasso, E. 2019. High heat tolerance in plants from the Andean highlands_: Implications for paramos in a warmer world. PLOS One, 10(14), e0224218

79: Liu, Y., Cao, T. & Glime, J. M. 2003. The Changes of Membrane Permeability of Mosses under High Temperature Stress. The Bryologist 106, 53-60 .

80: Logan BA, Monson RK. 1999. Thermotolerance of Leaf Discs from Four Isoprene-Emitting Species Is Not Enhanced by Exposure to Exogenous Isoprene. Plant Physiology 120: 821-826

81: Loik ME, Harte J. 1996. High-Temperature Tolerance of Artemesia tridentata and Potentilla gracilis. Oecologia 108: 224-231

82: Losch R. 1980. Die Hitzeresistenz der Pflanzen des kanarischen Lorbeerwaldes. Flora 170: 456-465

83: Maier R. 1971. Einflub von Photoperiode und Einstrahlungsst≈†rke auf die Temperaturresistenz einiger Samenpflanzen. ‚Ä¶sterreichische Botanische Zeitschrift, 119: 306-322

84: Marchin, RM, Backes, D, Ossola, A, Leishman, MR, Tjoelker, MG, Ellsworth, DS. 2021. Extreme heat increases stomatal conductance and drought-induced mortality risk in vulnerable plant species. Global Change Biology. 28: 1133-1146

85: Marchin, RM, Esperon-Rodriguez M, Tjoelker, MG, Ellsworth, DS. 2022. Crown dieback and mortality of urban trees linked to heatwaves during extreme drought. Science of the Total Environment 850: 157915

86: Marias, DE, Meinzer, FC, Woodru, DR, Mcculloh, KA. 2016. Thermotolerance and heat stress responses of Douglas- fi r and ponderosa pine seedling populations from contrasting climates. Tree Physiology, 301-315. https://doi.org/10.1093/treephys/tpw117

87: Methy, M, Gillon, D, Houssard, C. 1997. Temperature-induced changes of photosystem II activity in Quercus ilex and Pinus halepensis. Canadian Journal of Forest Research 27: 31-38

88: Meyer H, Santarius KA. 1998. Thermal Acclimation and Heat Tolerance of Gametophytes of Mosses. Oecologia 115: 1-8

89: Munchinger, IK, Hajek, P, Akdogan, B, Caicoya, AT, Kunert, N. 2023. Leaf thermal tolerance and sensitivity of temperate tree species are correlated with leaf physiological and functional drought resitance traits. Journal of Forestry Research 34: 63-76

90: Neuner G, Buchner O. 2013. Dynamics of Tissue Heat Toleranace and Thermotolerances of PS II in Alpine Plants (C. Lutz, Ed.).

91: Neuner, G, Buchner, O, Braun, V. 2000. Short-Term Changes in Heat Tolerance in the Alpine Cushion Plant Silene acaulis ssp. excapa [All.] J. Braun at Different Altitudes. Plant Biology 2:677-683

92: Neves Chaves, CJ, Santos Leal, BS, Lemos-Filho, JP. 2015. Temperature modulation of thermal tolerance of a CAM-tank bromeliad and the relationships with acid accumulation in different leaf regions. Physiologia Plantarum 154:500-510

93: Nobel PS. 1988. Environmental biology of agaves and cacti. New York, New York: Cambridge University Press.

94: Nobel, S, Geller, GN, Kee, SC, Departtnent, ADZ. 1986. Temperatures and thermal tolerances for cacti exposed to high temperatures near the soil surface. Plant Cell and Environment, 279-287

95: O'Sullivan OS, Heskel MA, Reich PB, Tjoelker MG, Weerasinghe KWLK, Penillard A, Zhu L, Egerton JJG, Bloomfield KJ, Creek D, et al. 2017. Thermal limits of leaf metabolism across biomes. Global Change Biology: 209-223

96: O'Sullivan OS, Weerasinghe KWLK, Evans JR, Egerton JJG, Tjoelker MG, Atkin OK. 2013. High-resolution temperature responses of leaf respiration in snow gum (Eucalyptus pauciflora) reveal high-temperature limits to respiratory function. Plant, Cell and Environment 36: 1268-1284

97: Offord CA. 2011. Pushed to the limit: Consequences of climate change for the Araucariaceae: A relictual rain forest family. Annals of Botany 108: 347-357

98: Okubo, N, Inoue, S, Ishii, HR. 2023. Tolerance and acclimation of the leaves of nine urban tree species to high temperatures. Forest. 14:1639

99: Ortiz, C, Cardemil, L. 2001. Heat-shock responses in two leguminous plants: a comparative study. J. Exp. Bot. 52: 1711-1719

100: Perez, TM, Feeley, KJ. 2020. Weak phylogenetic ad climatic constraints on photosynthetic heat tolerances. Journal of Biogeography. 48:91-100

101: Qiu, N, Lu, C. 2003. Enhanced tolerance of photosynthesis against high temperature damage in salt-adapted halophyte Atriplex centralasiatica plants. Plant, Cell and Environment 26:1137-1145

102: Ranney, TG, Ruter, JM. 1997. Foliar heat tolerance of three holly species (Ilex spp.): responses of chlorophyll fluorescence and leaf gas exchange to supraoptimal leaf temperatures. Journal of American Society of Horticultural Science. 122: 499-503

103: Reyes, M.A., Corcuera, LJ, Cardemil, L. 2003. Accumulation of HSP70 in Deschampsia antarctica Desv. leaves under thermal stress. Antarct. Sci. 15: 345-352.

104: Sapper I. 1935. Versuche zur hitzeresistenz der pflanzen. Planta 23: 518-556.

105: Sastry A, Barua D. 2017. Leaf thermotolerance in tropical trees from a seasonally dry climate varies along the slow-fast resource acquisition spectrum. Scientific Reports 7: 1-11.

106: Sastry, A, Guha, A, Barua, D. 2017. Leaf thermotolerance in dry tropical forest tree species: relationships with leaf traits and effects of drought. AOB Plants 10:plx070

107: Schreiber U, Berry JA. 1977. Heat-induced changes of chlorophyll fluorescence in intact leaves correlated with damage of the photosynthetic apparatus. Planta 136: 233-238

108: Seemann JR, Berry JA, Downton WJS. 1984. Photosynthetic Response and Adaptation to High Temperature in Desert Plants A Comparison of Gas Exchange and Fluorescence Methods for Studies of Thermal Tolerance. Plant Physiol 75: 364-368

109: Seemann JR, Downton WJS, Berry JA. 1986. Temperature and Leaf Osmotic Potential as Factors in the Acclimation of Photosynthesis to High Temperature in Desert Plants. Plant Physiology 80: 926-930

110: Singsaas L, Lerdau M, Winter K, Sharkey TD. 1997. lsoprene lncreases Thermotolerance of Isopreme-Emitting Species. Plant physiology: 1413-1420

111: Slot M, Krause GH, Krause B, Hernandez GG, Winter K. 2019. Photosynthetic heat tolerance of shade and sun leaves of three tropical tree species. Photosynthesis Research 141:119-130

112: Slot, M, Cala, D, Aranda, J, Virgo, A, Michaletz, ST, Winter, K. 2021. Leaf heat tolerance of 147 tropical forest species varies with elevation and leaf functional traits, but not with phylogeny. Plant Cell Environ. 44: 2414-2427

113: Smillie RM, Gibbons GC. 1981. Heat tolerance and heat hardening in crop plants measured by chlorophyll fluorescence. Carlsberg Research Communications 46: 395-403

114: Smillie RM, Nott R. 1979. Heat Injury in leaves of alpine, temperate and tropical plants. Australian Journal of Plant Physiology 6: 135-141

115: Smith, SD, Didden-Zopfy, B, Nobel, PS. 1984. High-Temperature Responses of North American Cacti. Ecology, 65: 643-651.

116: Snider, JL, Choinski, JS, Slaton, W. 2010. Juvenile leaves of Rhus glabra have higher photosynthetic thermal tolerance than mature leaves. Botany 88:286-289

117: Sonti, NF, Hallett, RA, Griffin, KL, Trammell, TLE, Sullivan, JH. 2021. Chlorophyll fluorescence parameters, leaf traits and foliar chemistry of white oak and red maple trees in urban forest patches. Tree Physiology 41: 269-679

118: Spiers, JA, Oatham, MR, Rostant, LV, Farrell, D. 2023. Determining the ecophysiological limits of a narrow niche tropical conifer tree (Podocarpus trinitensis) 43: 781-793

119: Srinivasam A, Hiroyuki T, Toshihiro S. 1996. Heat tolerances in food legumes as evaluated by cell membrane thermostability and chlorophyll fluorescence techniques. Euphytia 88: 35-45

120: Sumner, EE, Williamson, VG, Gleadow, RM, Wevill, T, Venn, SE. 2022. Acclimation to water stress improves heat tolerance to heat and freezing in a common alpine grass. Oecologia. 199:831-843

121: Tarvainen, L, Wittemann, M, Mujawamariya, M, Manishimwe, A, Zibera, E, Ntirugulirwa, B, Ract, C, Manzi, OJL, Andersson, MX, Spetea, C, Nsabimana, D, Wallin, G, Uddling, J. 2022. Handling the heat -- photosynthetic thermal stress in tropical trees. New Phytologist 233(1):236-250

122: Terzaghi WB, Fork DC, Berry JA, Field CB. 1989. Low and High Temperature Limits. Plant Physiology: 1494-1500

123: Tiwari, R, Gloor, E, Jonatar, W, Cruz, A, Marimon, BS, Marimon-junior, BH, √â Vitria, AP. 2020. Photosynthetic quantum efficiency in south-eastern Amazonian trees may be already affected by climate change. Plant Cell and Environment 1-12. https://doi.org/10.1111/pce.13770

124: Valladares F, Pearcy RW. 1997. Interactions between water stress, sun-shade acclimation, heat tolerance and photoinhibition in the sclerophyll Heteromeles arbutifolia. Plant, Cell And Environment 20: 25-36

125: Valliere, JM, Nelson, KC, MCastaneda Martinez, M. 2023. Functional traits and drought strategy predict leaf thermal tolerance. Conservation Physiology 11(1): coad085

126: Visakorpi, K, Manzanedo, RD, Gorlich, AS, Schiendorfer, K, AltermanttBieger, A, Gates, E, HilleRisLambers, J. 2023. Leaf-level resistance to frost, drought, and heat covaries across European temperate tree seedlings. Journal of Ecology 112: 559‚Äì574

127: Weng J, Lai M. 2005. Estimating heat tolerance among plant species by two chlorophyll fluorescence parameters. Photosynthetica 43: 439-444

128: Zhang JL, Poorter L, Hao GY, Cao KF. 2012. Photosynthetic thermotolerance of woody savanna species in China is correlated with leaf life span. Annals of Botany 110: 1027-1033

129: Zhu L, Bloomfield KJ, Hocart CH, Egerton JJG, O?Sullivan OS, Penillard A, Weerasinghe LK, Atkin OK. 2018. Plasticity of photosynthetic heat tolerance in plants adapted to thermally contrasting biomes. Plant Cell and Environment 41: 1251-1262

130: Zohar, Y, Waisel, Y, Karschon, R. 1981. Heat and cold resistance of Eucalyptus occidentalis Endl. Leaves and its relationship to soil water conditions. Australia Journal of Ecology 6:79-84

131: Zsofi Z, Varadi G, Balo B, Marschall M, Nagy Z, Dulai S. 2009. Heat acclimation of grapevine leaf photosynthesis: Mezo- and macroclimatic aspects. Functional Plant Biology 36: 310-322

**Section 3: IPCC Region heat tolerances**

Table S5: The mean heat tolerances for IPCC regions.

| Region | Tcrit | T50 | Tmax |  | Region | Tcrit | T50 | Tmax |
| --- | --- | --- | --- | --- | --- | --- | --- | --- |
| Arabian-Peninsula | NA | NA | NA |  | N.Pacific-Ocean | NA | NA | NA |
| Arabian-Sea | NA | NA | NA |  | N.South-America | 50.3 | NA | NA |
| Arctic-Ocean | NA | NA | NA |  | N.W.North-America | 41.8 | NA | NA |
| Bay-of-Bengal | NA | NA | NA |  | N.W.South-America | 48.3 | 49.2 | 56.2 |
| C.Australia | 53.8 | NA | NA |  | New-Zealand | 48.4 | NA | NA |
| C.North-America | 44.8 | NA | NA |  | Russian-Arctic | NA | NA | NA |
| Caribbean | NA | 47.9 | NA |  | Russian-Far-East | NA | NA | NA |
| Central-Africa | NA | NA | NA |  | S.Asia | 45.5 | 48.8 | NA |
| E.Antarctica | NA | NA | NA |  | S.Atlantic-Ocean | NA | NA | NA |
| E.Asia | NA | 44.7 | NA |  | S.Australia | 47.2 | NA | NA |
| E.Australia | 44 | 50.5 | 56.9 |  | S.Central-America | 46.9 | 50.1 | 52.8 |
| E.C.Asia | NA | NA | NA |  | S.E.Asia | NA | NA | NA |
| E.Europe | NA | NA | NA |  | S.E.South-America | NA | NA | NA |
| E.North-America | 47.4 | NA | NA |  | S.Eastern-Africa | NA | NA | NA |
| E.Siberia | NA | NA | NA |  | S.Indic-Ocean | NA | NA | NA |
| E.Southern-Africa | NA | NA | NA |  | S.Pacific-Ocean | 46.5 | 52.7 | 58.5 |
| Equatorial.Atlantic-Ocean | NA | NA | NA |  | S.South-America | NA | NA | NA |
| Equatorial.Indic-Ocean | NA | NA | NA |  | S.W.South-America | NA | NA | NA |
| Equatorial.Pacific-Ocean | NA | NA | NA |  | Sahara | NA | 49.2 | NA |
| Greenland/Iceland | 48.2 | NA | NA |  | South-American-Monsoon | 43.1 | 52.3 | 62.4 |
| Madagascar | NA | NA | NA |  | Southern-Ocean | NA | NA | NA |
| Mediterranean | NA | 50.5 | NA |  | Tibetan-Plateau | NA | NA | NA |
| N.Atlantic-Ocean | NA | NA | NA |  | W.Antarctica | NA | NA | NA |
| N.Australia | 46.2 | 52.2 | 58.3 |  | W.C.Asia | NA | NA | NA |
| N.Central-America | 53.4 | NA | NA |  | W.North-America | 47.7 | 47.6 | 56.8 |
| N.E.North-America | NA | NA | NA |  | W.Siberia | NA | NA | NA |
| N.E.South-America | 46.9 | 47.7 | NA |  | W.Southern-Africa | NA | NA | NA |
| N.Eastern-Africa | NA | NA | NA |  | West&Central-Europe | 49.4 | 51 | 53.1 |
| N.Europe | 47.2 | 47.8 | NA |  | Western-Africa | NA | 48.5 | NA |

We mapped occurrences to their growth site coordinates if they also has provenance data to estimate the mean heat tolerance of each IPCC region. This was done to highlight records with a full compliment of biogeographical data. We then calculated the mean heat tolerance regardless of method for all records within a given region. The mean Tcrit for IPCC regions ranged from 41.8˚C in NW North America to 53.8 in Central Australia, from 47.6˚C in W. North America to 52.7 in the Southern pacific Ocean for T50, and from 52.8 in S. Central America to 62.4 in the South American Monsoon region for Tmax.
